# Cytosolic DNA structures produced by mismatch-repair deficiency coordinate anti-tumor immunity in colorectal cancer

**DOI:** 10.1101/2025.07.17.665440

**Authors:** Shayla R. Mosley, Natalie Lapa, Afshin Namdar, Thomas E. Catley, Felix Meier-Stephenson, Courtney Mowat, Pinzhang Gao, Xuejun Sun, Vanessa Meier-Stephenson, Alice L. B. Pyne, Kristi Baker

## Abstract

Patients with the microsatellite instable (MSI) subtype of colorectal cancer (CRC) have better prognosis and immunotherapy response than patients with the chromosomally instable (CIN) subtype due to improved cytotoxic T cell responses from high neoantigen levels and production of the chemokines CXCL10 and CCL5 that recruit cytotoxic T cells. This high chemokine production in MSI CRCs is due to constitutive activation of the cytosolic DNA (cyDNA) sensor STING by specific features of MSI cyDNA that lead to more effective STING pathway activation. Here, we investigate the features of MSI and CIN cyDNA to identify structures that more effectively activate STING. We find that MSI cyDNA is enriched in G-quadruplexes which improve STING and CD8^+^ T cell activation. Additionally, MSI micronuclei are also more efficient at inducing chemokine expression than CIN micronuclei. However, micronuclei are less effective than free cyDNA at inducing anti-tumor immunity and instead lead to increased Treg activation and IL10 production. Overall, these data highlight the role of specific cyDNA structures in anti-tumor immunity and provide essential knowledge for improved design of therapeutic DNA-based STING agonists that could be combined with immune checkpoint therapies to improve the prognosis of poorly immunogenic tumors like CIN CRCs.

## Introduction

Colorectal cancer (CRC) is the second most deadly cancer worldwide, with an estimated 1.9 million people diagnosed per year^1^. The majority of CRC cases are chromosomally instable (CIN), arising from dysregulated WNT signaling through loss of *APC* and further accumulation of other oncogenic driver mutations in *KRAS*, *BRAF*, *PTEN*, and *TP53*^2,3^. However, 12-15% of CRC tumors instead have microsatellite instability (MSI) arising due to deficient mismatch repair due to epigenetic silencing of *MLH1*^4^. MSI CRCs therefore have widespread point mutations and frameshifts especially in microsatellite regions of the genome^4^. Clinically, patients with MSI CRC have a 35% lower risk of death^5^ due to improved cytotoxic T cell infiltration and activation than CIN tumors^6^. Consistent with this, MSI patients treated with the checkpoint inhibitor therapy pembrolizumab have an objective response rate of 40%, compared to 0% of CIN CRC patients^7^. This is partially due to higher neoantigen production in MSI CRCs that arise from the hypermutability of MSI tumors and lead to greater cytotoxic T cell recognition^4^. However this explanation is incomplete as it fails to explain why MSI tumors in other organs lack improved CD8^+^ T cell responses^8^, highlighting the contribution of neoantigen independent mechanisms in MSI CRC immunity. We previously showed that high production of the chemokines CXCL10 and CCL5 is essential for CD8^+^ T cell infiltration into MSI CRCs, an effect found to be dependent on activation of the STING pathway^9^.

STING is a pattern recognition receptor (PRR) that responds to microbial DNA in the cytosol^10^. This DNA is sensed and directly bound by cGAS, which produces the cyclic dinucleotide 2’3’-cGAMP, a process that is improved by oligomerization of multiple cGAS proteins with DNA^11,12^. 2’3’-cGAMP binds STING, inducing STING dimerization to act as a scaffold for the phosphorylation of TBK1, STING, and the transcription factor IRF3 by TBK1^13^. IRF3 then translocates to the nucleus to induce transcription of type I interferons (IFN) IFNα and IFNβ, which then trigger expression of interferon stimulated genes (ISGs) such as CXCL10 and CCL5 through the JAK/STAT pathway^12,14^. In recent years, cGAS/STING have also emerged as key monitors of genome integrity, as DNA damage from genomic instability or exogenous sources causes a cell’s own DNA to leak into the cytosol as endogenous cytosolic DNA (cyDNA)^14,15^. This cyDNA can be found in two forms: small (typically <200bp) free DNA fragments or micronuclei where larger chromosomes or chromosome fragments are encased in an often unstable nuclear envelope^14–16^. CyDNA has important implications in cancer, where STING plays a role in cancer prevention and effective response to DNA damaging therapies such as radiation (IR)^17,18^. In line with this, STING agonists are currently in clinical trials in combination with checkpoint inhibition therapies such as pembrolizumab^19,20^. Importantly, STING has also been shown in certain contexts to induce pro-tumorigenic effects^21–23^, however how STING is induced to act in a pro- vs anti-tumorigenic manner is not fully understood.

Dendritic cells (DCs) are important regulators of the transition from innate to adaptive immunity, and are key mediators of STING responses^12^. DCs are activated by exogenous type I IFN, leading to improved antigen presentation and co-stimulation of CD8^+^ T cells^12^. Importantly, DCs have recently been shown to uptake 2’3’-cGAMP and cyDNA from tumor cells through gap junctions, phagocytosis, and exosomes, leading to activation of DC intrinsic STING and type I IFN production as well as greater CD8^+^ T cell activation^12,24,25^. We previously demonstrated the critical role of DCs in MSI vs CIN CRC immunity where uptake of highly stimulatory cyDNA patterns induced a type I IFN gene signature, including CXCL10 and CCL5, and improved CD8^+^ T cell infiltration and activation in CIN tumors^16^.

Critically, the intensity of this response varied according to specific features present in MSI cyDNA such as microsatellite repeats, which we showed increased STING activation and thereby CD8^+^ T cell activation at equal concentrations^16^. Additionally, DNA damaging therapies could influence the stimulatory nature of cyDNA, with IR producing cyDNA more stimulatory to STING in part due to increased cyDNA size and mitochondrial DNA content^16^. However, a key unresolved question in the field is the role of secondary or tertiary DNA structure on STING activation, which are not only frequently formed by microsatellite sequences^26^ but are also highly likely to influence cGAS-DNA binding and oligomerization. Additionally, these works largely focused on free cyDNA and failed to evaluate the role of micronuclei in MSI and CIN CRCs. Indeed, much of the current literature focuses solely on either free or micronuclear cyDNA alone or fails to separate the distinct roles these cyDNA types may have on STING activation. This is critical to understand given the frequent association of chromosomal instability with micronuclei in CRCs and other tumor types^27,28^.

Here, we investigate the role of cyDNA structures in MSI and CIN cyDNA, particularly in terms of G-quadruplexes (G4s), a DNA structure formed by guanine repeats common within promoter regions that influence gene transcription^29^. We find G4s are heavily represented in MSI cyDNA and are capable of strong STING activation. This is especially important as G4 binding agents are currently under investigation for cancer treatment^30^, and we find these agents also produce cyDNA more effective at activating STING. We show that MSI micronuclei are also more effective at activating STING than CIN micronuclei, consistent with our previous findings for free MSI cyDNA^16^. However, our findings indicate that micronuclei are less capable of inducing anti-tumor CD8^+^ T cell activation than free cyDNA, and instead lead to higher Treg activation and expression of the immuno-inhibitory cytokine IL10. This insight sheds light on a key immune inhibitory mechanism that has previously been unrecognized in CIN CRCs. Overall, we show that different structural confirmations within endogenous cyDNA can be used to activate or inhibit CD8^+^ T cell responses, providing further insight into the design of DNA based STING agonists or STING activating therapies aimed to promote anti-tumor immune responses in immunologically cold tumors.

## Results

### CyDNA contains various DNA structures

To investigate the structure of CRC cyDNA, we isolated cyDNA from MC38 cells rendered MSI or CIN by CRISPR induced mutation^9^, separated the cyDNA to similar sizes by size exclusion chromatography (Figure S1A), and performed circular dichroism spectroscopy. Both large (>45 bp) and small (∼40 bp) cyDNA fragments exhibited a broad positive band at approximately 260nm-280nm with a negative band at 245nm (Figure 1A), suggesting cyDNA is largely composed of structured B-DNA, also known as standard right-handed double helix DNA^31^.

**Figure 1.**
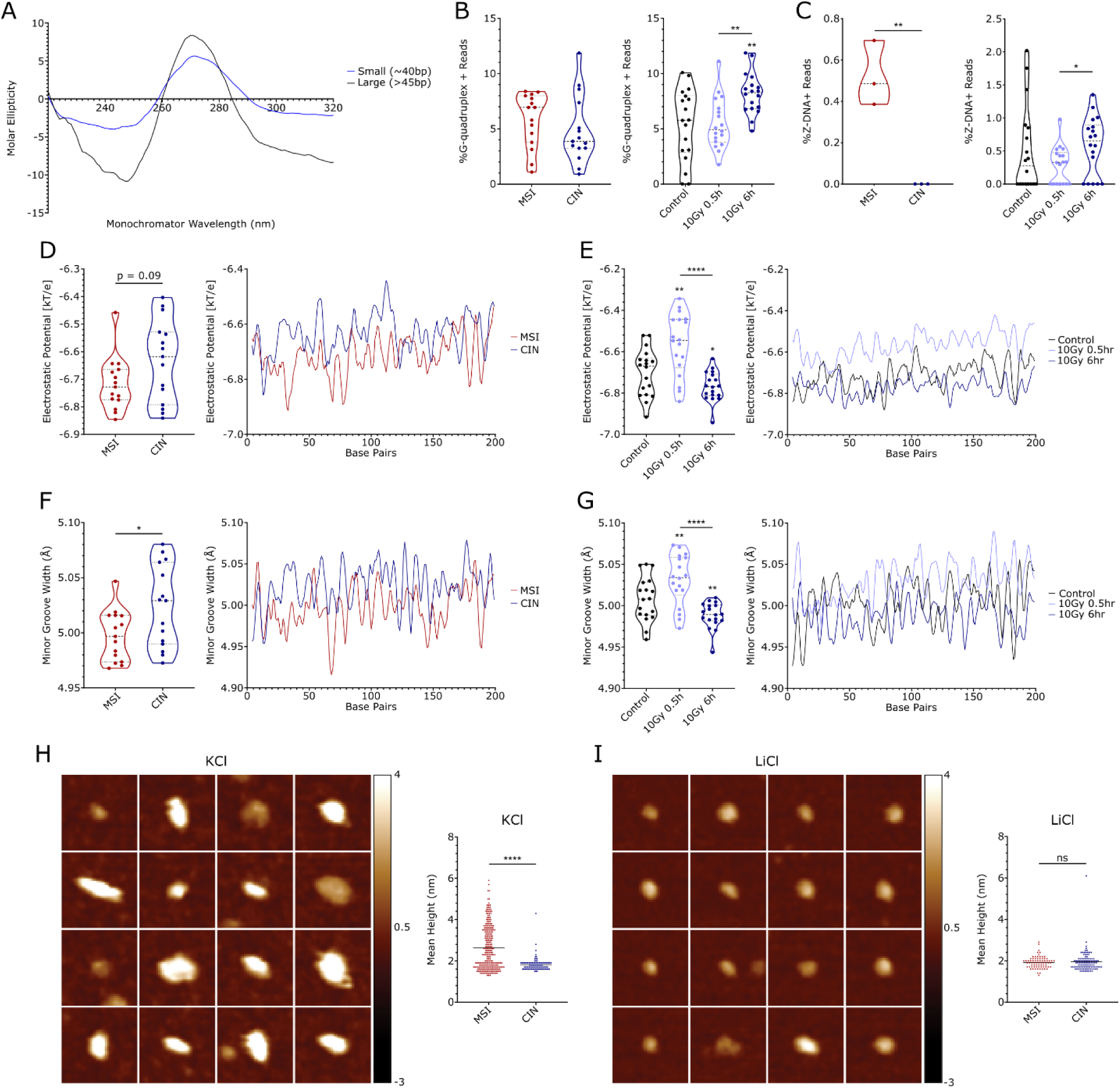
CyDNA contains various secondary structures. (A) CyDNA isolated from MSI CRC cells were separated by size exclusion chromatography before analysis of cyDNA structure by circular dichroism. Representative replicate of n = 2. (B-G) MSI and CIN CRC cells were treated with 10Gy IR (0.5 or 6 hours of recovery), the indicated concentration of 5-FU for 24 hours, or left untreated before isolation of cyDNA and next generation sequencing, n = 3. (B) G4Hunter analysis of G4 sequence prevalence in MSI vs CIN cyDNA (all treatments combined, left) or in cyDNA following IR treatment (all cell types combined, right). (C) Percentage of predicted Z-DNA sequences in MSI vs CIN (untreated, left) or IR treated cyDNA (all cell types combined, right). (D-G) DNAShapeR analysis of electrostatic potential (D-E) and minor groove width (F-G) in MSI vs CIN (all treatment types combined) or IR treated (all CRC subtypes combined) cyDNA. Average read values are indicated on the left for each sample, average mean per base for each treatment group is indicated on the right. (H-I) MSI and CIN cyDNA were separated using size exclusion chromatography before AFM analysis in KCl (H) or LiCl (I) buffers. (Left) Representative AFM images of small (∼40bp) MSI cyDNA fragments. Each tile is 40×40nm, z-scale bar of height is -3 to 4nm. (Right) Mean height per particle of small (∼40bp) MSI or CIN cyDNAs. ns indicates non-statistically significant, >70 DNA strands were imaged per sample. Statistics were determined by paired (B, C right, D-G) or unpaired (C left, H, I) T-test, * = p < 0.05, ** = p < 0.01, **** = p < 0.0001.

To determine if other DNA structures were present in cyDNA at lower concentrations, we sequenced cyDNA isolated from MSI or CIN MC38 cells that had been treated or not with the DNA damaging therapies IR or 5-fluorouracil (5-FU)^16^. Analysis of these reads with G4Hunter, a program to quantify sequences capable of forming G4s^32^, indicated a trending increase in G4s in MSI cyDNA compared to CIN (Figure 1B). IR also led to increased predicted G4s in cyDNA, but 5-FU did not (Figure S1B). Conversely, analysis of G4 content by the Gquad package, which predicts multiple DNA structures including G4s, Z-DNA, triplexes, as well as repetitive sequences, showed no increase in predicted G4 content in the cytosol of MSI or IR treated cells (Figure S1C). Z-DNA is an alternative DNA structure that forms a left-handed helix with a zig zag backbone and has previously been implicated in poor cGAS activation^33^. Z-DNA motifs were also increased in untreated MSI cyDNA compared to CIN, and were further induced by IR treatment (Figure 1C) but not by treatment with 5-FU (Figure S1D). IR also led to increased levels of predicted triplexes in CRC cyDNA (Figure S1E). Consistent with our previous findings of increased microsatellites in MSI cyDNA^16^, more short tandem repeats and slipped motifs were found in MSI cyDNA compared to CIN cyDNA (Figure S1F-G). IR, but not 5-FU, also increased short tandem repeat and slipped motifs in CRC cyDNA. Overall, CRC cyDNA contains many differences in predicted structure, which may include increased G4s in MSI cyDNA.

To assess other structural components of cyDNA that could influence cGAS binding, we utilized DNAShapeR, a package that quantifies DNA properties known to influence transcription factor binding^34^. We specifically focused on electrostatic potential and minor groove width, as cGAS binds the minor groove of DNA through electrostatic interactions^11,35,36^. MSI cyDNA and cyDNA arising from IR treatment contained sequences with lower electrostatic potential (Figure 1D-E, S2A-D). In contrast, 5-FU led to cyDNA with a more positive electrostatic potential (Figure S2E-G). MSI cyDNA and IR treated cyDNA, but not 5-FU treated cyDNA, also contained sequences indicative of smaller minor groove width (Figure 1F-G, S3). Therefore, highly stimulatory MSI and IR-treated cyDNAs exhibit tighter minor grooves and lower electrostatic potential.

Notably, prediction of cyDNA structure by sequence only indicates that the sequences needed to form a structure are present, not whether the structure actually occurs since this is dependent on other factors such as appropriate ionic conditions. We therefore sought to examine structure using a more direct method. Atomic force microscopy (AFM) has previously been shown to be highly effective at detection of structures such as G4s^37^. We thus separated MSI and CIN cyDNA by size using size exclusion chromatography and evaluated the structure of cyDNA by AFM. Results showed that the majority of large (>45bp) and small (∼40bp) cyDNA fragments are 1-2nm in height (Figure 1H, S4), indicative of ss and dsDNA respectively^37^. Additionally, cyDNA isolated from MSI cells exhibited DNA fragments 3-5nm in height, suggesting structures containing 3-5 DNA strands were present. As this is consistent with 4 stranded G4 structures, we repeated AFM imaging in the presence of LiCl, which is known to destabilize G4s^38^. Addition of LiCl effectively removed the 3-5nm DNA fragments (Figure 1I, S4), suggesting that MSI cyDNA is enriched for G4s.

### G4s in MSI cyDNA increase STING activation

G4s are tertiary DNA structures resulting from guanine rich regions (at minimum G≥_3_N_x_G≥_3_N_x_G≥_3_N_x_G≥_3_) that form Hoogsteen hydrogen bonds between guanine residues to create a tower-like quadruplex structure^29^. These G4 structures may contain many variations, such as in size by the number of bonded guanines, the parallel, antiparallel, or hybrid structures depending on the orientation of the connecting linker strands, or the presence of bulges due to non-continuous guanine repeats^29^. G4 sequences are frequently found in promoter regions and telomeres^29^ and are enriched in many viral genomes^39^. Therefore, to determine if the presence of G4s in MSI cyDNA helped to explain its strong ability to activate STING, G4s were pulled down from the cytosol of MSI or CIN cells and equal concentrations were used to stimulate bone marrow derived dendritic cells (BMDCs) (Figure 2A). Equal concentrations of G4 enriched cyDNA from MSI or CIN cells led to strong induction of *Cxcl10* and *Ccl5* compared to G4 depleted cyDNA or total cyDNA (Figure 2B), suggesting G4s are potent inducers of STING. Of note, G4 enriched cyDNA also led to equal or higher *Cxcl10* and *Ccl5* expression to herring testis DNA (HT-DNA), a STING agonist commonly used as a positive control in the literature. To confirm these results were not a due to the pulldown process itself, we repeated the stimulation with oligos containing known G4 forming sequences from promoter regions (Table S1). Oligos A and B contained G4 sequences from the *HRAS* promoter that are known to form parallel and anti-parallel G4s respectively^40^, while oligo C contained the sequence for the antiparallel G4 in the *MDM2* promoter^41^. DNA oligos were induced to form structure by heating to 95°C and slow cooling overnight before separation of oligos that formed structure (hereafter deemed G4 high) from oligos that failed to form structure (deemed G4 low) by size exclusion chromatography (Figure S5A). Notably, separation by this method ensured that any observed differences would not be due to the presence of highly stimulatory repetitive unstructured sequences. To confirm G4s were formed and effectively separated by this process, G4 high and G4 low fractions of oligo C were visualized by circular dichroism spectroscopy. The G4 high oligos exhibited a negative peak at approximately 250nm and a broad positive peak from approximately 270-310nm (Figure S5B), consistent with previous reports of the *MDM2* anti-parallel G4^41^. To evaluate the ability of G4 high vs G4 low oligos to induce STING activation, equal concentrations were used to stimulate BMDCs and STING activation was determined by RNA isolation and RT-qPCR. Interestingly, the G4 high oligo C led to increased expression of *Irf7* compared to the G4 low oligo of the same sequence, although no consistent differences were observed between the G4 high and low oligos A and B (Figure 2C). Next, to determine if G4s led to increased co-stimulation of CD8^+^ T cells by DCs, BMDCs were stimulated with G4 high or low oligos in the presence of the model neoantigen ovalbumin (OVA) for 30 minutes. These stimuli were then washed away before addition of OT-1 CD8^+^ T cells for 48 hours and evaluation of T cell activation by flow cytometry. Compared to the respective G4 low oligos, G4 high oligo B led to increased Ki67 and CD69, while G4 high oligos B and C increased CCR5 on CD8^+^ T cells (Figure 2D), indicating increased T cell proliferation, activation, and response to IFN respectively. No consistent differences in CD8^+^ T cell activation were observed between G4 high and low oligos A, suggesting that only certain G4 structures are more effective at inducing STING or CD8^+^ T cell activation compared to their unstructured sequence.

**Figure 2.**
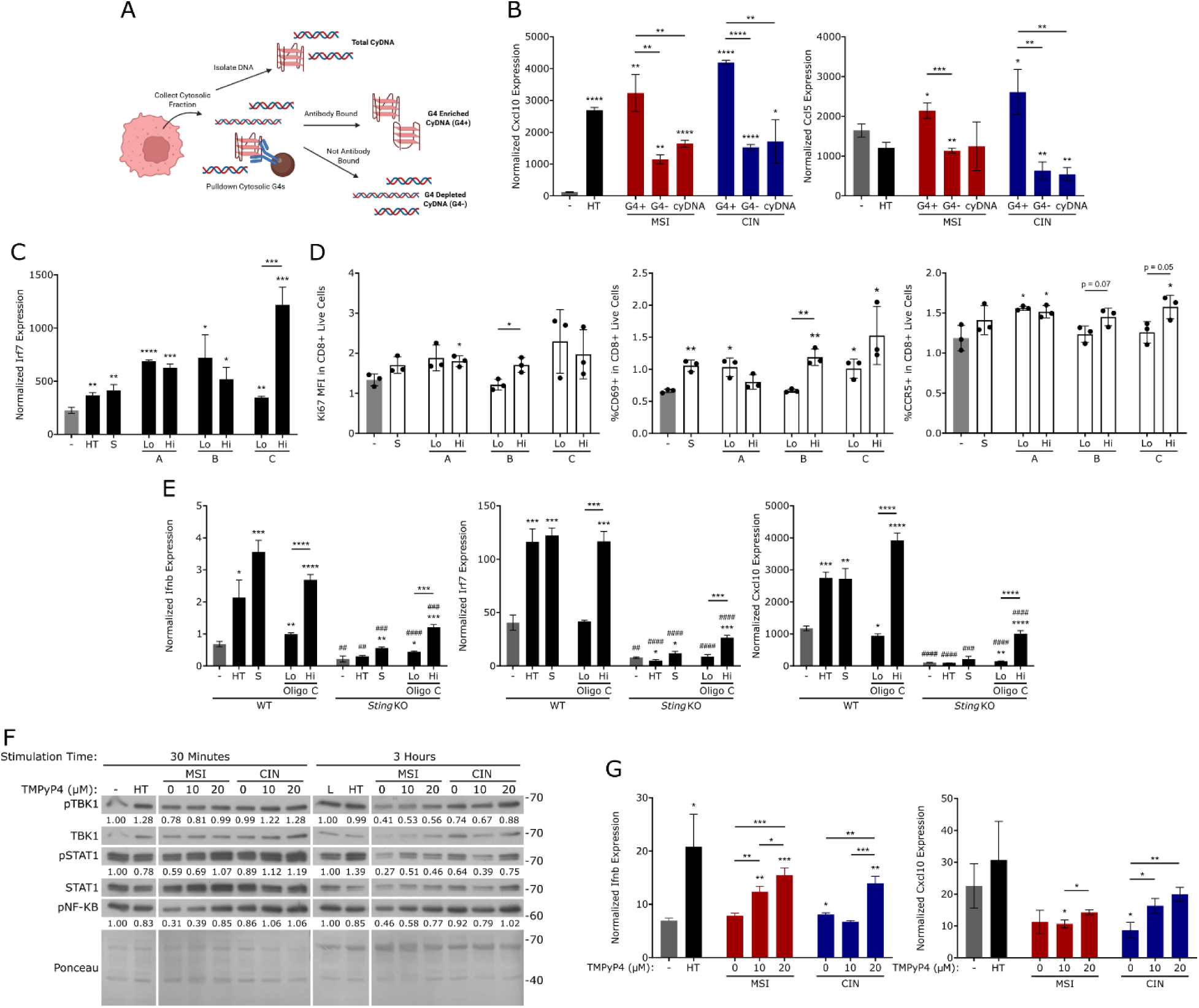
G4s in MSI cyDNA increase STING and CD8^+^ T cell activation. (A-B) G4s were pulled down from the cytosol of MSI and CIN cells and 500ng was used to stimulate BMDCs with lipofectamine for 4 hours before isolation of RNA and RT-qPCR. G4+ indicates cyDNA pulled down with a G4 antibody, G4-indicates cyDNA remaining after pulldown with a G4 antibody, cyDNA indicates total cyDNA, HT indicates herring testis DNA, - indicates the lipofectamine vehicle control. Representative replicate shown of n = 3. (C) G4 high (Hi) and G4 low (Lo) oligos of sequence A, B, or C were used to stimulate BMDCs for 3 hours before isolation of RNA and RT-qPCR. S indicates double stranded scramble control. Representative replicate shown of n = 3. (D) BMDCs were stimulated with G4 high (Hi) and G4 low (Lo) oligos and OVA protein for 30 minutes before washing and co-culture with OT-1 CD8^+^ T cells for 48 hours and evaluation of CD8^+^ T cell activation (CD69), proliferation (Ki67), and response to IFN (CCR5) by flow cytometry. Representative replicate shown of n = 3 with 3 biological replicates each. Data shown is normalized to T cell only controls. (E) *Sting* knockout (KO) BMDCs or wildtype (WT) BMDCs were stimulated with 500ng of G4 high (Hi) or G4 low (Lo) oligo C for 3 hours before isolation of RNA and RT-qPCR. Representative replicate shown of n = 2. # indicates statistical significance to the same stimulus on WT BMDCs. (F-G) MSI and CIN cells were treated for 48 hours with the indicated concentration of the G4 binder TMPyP4 before isolation of cyDNA. 500ng TMPyP4 induced cyDNA was then used to stimulate BMDCs for 30 minutes before removal of stimulus, washing, and incubation up to the total stimulation time of 30 minutes or 3 hours for collection of protein (F) or 24 hours for collection of RNA (G). Representative replicate shown of n = 2. Quantifications shown are normalized to the Ponceau loading control and lipofectamine vehicle control. Protein ladder sizes are shown in kDa. * over the sample bar indicates statistical significance compared to the vehicle control. Statistical significance was determined by unpaired T test, * or # = p < 0.05, ** or ## = p < 0.01, *** or ### = p < 0.001, **** or #### = p < 0.0001.

Next, to confirm that G4s were increasing cytokine expression through STING and not another PRR, BMDCs isolated from *Sting^Gt/Gt^* knockout or wild type mice were stimulated with G4 high and low oligo C. *Sting* knockout BMDCs induced substantially less expression of *Ifnβ*, *Irf7* and *Cxcl10* in response to G4 high oligo C compared to wildtype BMDCs, although G4 high oligo C still induced higher expression of these genes than G4 low oligo C in *Sting* knockout cells (Figure 2E), indicating that G4s lead to more effective STING activation, but may also induce activation of other PRRs. Interestingly, we also did not observe better binding of cGAS to G4 high vs G4 low oligos by electrophoretic mobility shift assay (EMSA) (Figure S5C), suggesting G4 high oligos B and C increase STING and CD8^+^ T cell activation by a mechanism other than simple binding affinity.

Finally, we sought to determine if endogenous cytosolic G4 content could be increased as a potential therapeutic approach to induce improved STING activation in tumors. G4 binding compounds are currently undergoing clinical trials for use against cancer and act by inhibiting telomerase and oncogene transcription, as well as inducing genomic instability and DNA damage^30,42^. As DNA damage by these compounds is often due to replication fork stalling and collapse at G4s^30^, we reasoned that DNA damage at these sites would increase cytosolic G4 content. Additionally, introduction of cytosolic G4s by this method would likely lead to a mixture of different G4 structures, which our stimulation data (Figure 2B-C) suggests may be a more effective method of increasing STING activation. TMPyP4 is a G4 stabilizing ligand previously shown to increase DC and CD8^+^ T cell infiltration into tumors through activation of STING by increasing genomic G4s and double strand breaks^43^. To determine if TMPyP4 treatment led to cyDNA more capable of activating STING, MSI and CIN cells were treated for 48 hours with TMPyP4 before isolation of cyDNA. Equal concentrations of this cyDNA were then used to stimulate BMDCs to examine STING activation. CyDNA from TMPyP4 treated cells led to increased phosphorylation of TBK1, STAT1, and NF-κB, as well as expression of *Ifnβ* and *Cxcl10* (Figure 2F-G), indicating TMPyP4 treatment successfully increases the ability of endogenous cyDNA to activate STING at equal DNA concentrations.

### MSI micronuclei are more efficient at activating PRRs than CIN micronuclei

While our previous data indicated cyDNA from MSI CRCs was more efficient at inducing STING activation and thereby more effective at inducing CD8^+^ T cell activation by DCs^16^, these investigations and those above largely focused on free fragmented cyDNA and failed to evaluate the role of micronuclear DNA in this process. This is an important knowledge gap to address given the highly structured nature of micronuclear DNA and its known association with CIN^27,28^. Therefore, to evaluate the role of micronuclei in STING activation in MSI and CIN CRCs, we first quantified micronuclei content in these cells by immunofluorescence microscopy (Figure S6A). Consistent with previous reports^44^, both MSI and CIN CRC cells showed highly increased levels of micronuclei 24 hours following IR treatment (Figure 3A). However, we surprisingly observed no significant difference between the number of micronuclei in MSI and CIN cells, suggesting the number of micronuclei does not influence differences in STING activation between MSI and CIN CRCs.

**Figure 3.**
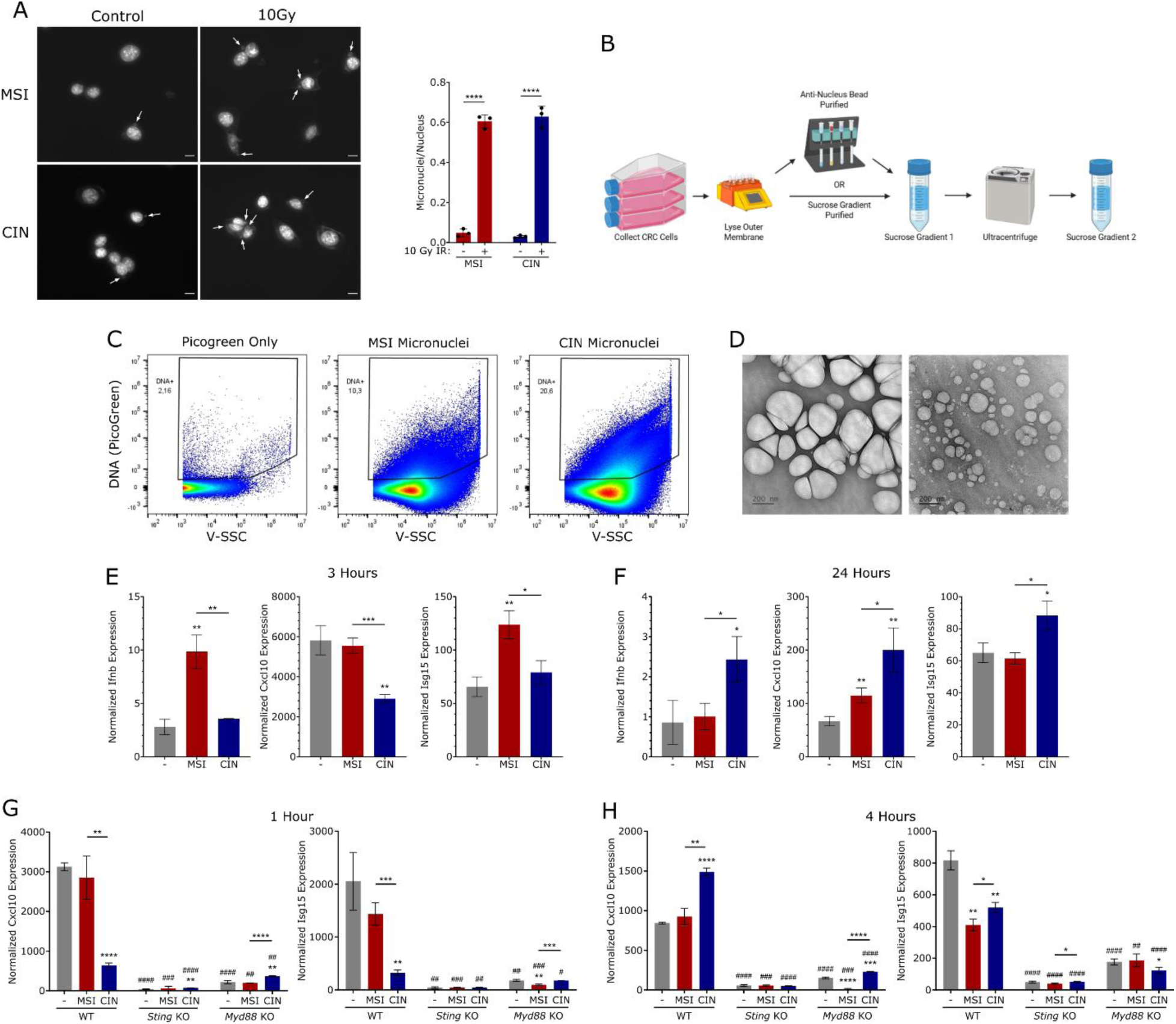
MSI micronuclei are more efficient at inducing PRR activation at equal DNA concentrations. (A) MSI and CIN cells were left untreated or were treated with 10Gy IR and allowed to recover for 24 hours before fixation and quantification of micronuclei by fluorescence imaging of DAPI stained cells. (Left) Representative images with arrows indicating micronuclei, size bar indicates 10µm. (Right) Quantification of micronuclei in the fluorescent images. Representative replicate shown of n = 3 with 3 biological replicates each and >100 cells per biological replicate. (B) Schematic of micronuclei isolation made in BioRender. CRC cells were collected by Trypsinization before lysis of the outer membrane on the gentleMACS Dissociator. Lysates then either proceeded to the next step or were purified using anti-nucleus microbeads before proceeding to the next step (bead purified micronuclei). Organelles were then separated by density and size using 2 sucrose gradients. (C) Microparticle flow cytometry of micronuclei isolated from MSI and CIN cells and stained with the DNA stain PicoGreen. Representative replicate of n = 3. (D) Transmission electron microscopy images of micronuclei isolated from MSI cells, size bars shown are 200nm. Representative replicate shown of n = 2. (E-F) Micronuclei were isolated from MSI and CIN cells and 100ng was used to stimulate BMDCs for 3 (E) or 24 hours (F) before RNA isolation and RT-qPCR. – indicates PBS vehicle control. (G-H) Micronuclei isolated from MSI and CIN cells were used to stimulate wildtype (WT), *Sting^Gt/Gt^* (*Sting* KO), or *Myd88* knockout (*Myd88* KO) BMDCs for 1 (G) or 4 hours (H) before isolation of RNA and RT-qPCR. Representative replicate shown of n = 2. Statistical significance was determined by unpaired T test. * over the sample bar indicates significance to the vehicle control, # over the sample bar indicates significance to the same condition with wildtype BMDCs. * or # = p < 0.05, ** or ## = p < 0.01, *** or ### = p < 0.001, **** or #### = p < 0.0001.

Next, to examine the ability of MSI and CIN micronuclei to activate STING, we isolated micronuclei from MSI and CIN CRC cells by cell fractionation and subsequent sucrose gradient separation (Figure 3B). These isolated particles contained a high percentage of DNA^+^ particles as assessed by microparticle flow cytometry (Figure 3C) and were spherical in shape and heterogeneous in size as assessed by electron microscopy (Figure 3D). Notably, we did not detect mitochondrial contamination by COX-IV protein content (Figure S6B) or observe rod-shaped particles by electron microscopy (Figure 3D), confirming our successful isolation of pure micronuclei. Additionally, no differences were observed in the levels of cGAS present in MSI and CIN micronuclei (Figure S6C). To examine STING activation in DCs by these micronuclei, equal amounts of micronuclei (as determined by DNA content analysis via PicoGreen dsDNA stain) were used to stimulate BMDCs before analysis of their gene expression. Consistent with previous observations with free cyDNA^16^, MSI micronuclei induced increased levels of *Ifnβ*, *Cxcl10*, and *Isg15* transcripts at 3 hours compared to CIN micronuclei (Figure 3E). However, by 24 hours, CIN micronuclei induced stronger expression of these transcripts (Figure 3F). Next, to determine if micronuclei-induced cytokine expression is through the STING pathway or through other PRRs, micronuclei isolated from MSI and CIN cells were used to stimulate BMDCs with knockout of *Sting* or *Myd88*, the latter being a key effector in the TLR pathway^10^. *Cxcl10* and *Isg15* transcription induced by both early (MSI) and late (CIN) micronuclei responses were diminished with loss of *Sting* or *Myd88* (Figure 3G-H), suggesting communication through multiple PRR pathways, including STING and TLRs, is required for effective response to micronuclei. Therefore, MSI micronuclei are more efficient at activating PRRs such as STING than CIN micronuclei.

One caveat of these experiments is that micronuclei purification by sucrose gradient is solely based on size and density. This method is current standard practice and is beneficial since more targeted isolation methods may bias for micronuclei of certain protein or membrane composition. However, this method can lead to contamination from other intracellular organelles which may have unknown effects on DC activation. Although we observed very little signs of such contamination (Figures 3C-D, S6B), we sought a second method of micronuclei isolation to confirm our findings. Therefore, we utilized anti-nucleus microbeads which target nuclear membrane molecules to further purify our micronuclei isolations (Figure 3B). These bead purified micronuclei also contained high levels of DNA^+^ particles (Figure S6D), no detectable mitochondrial contamination (Figure S6B), and were spherical in shape (Figure S6E) consistent with those isolated solely by density centrifugation. Consistent with our previous data (Figure 3E-F), stimulation of BMDCs with equal DNA concentrations of bead purified micronuclei led to increased *Cxcl10* and *Ccl5* expression at early time points from MSI micronuclei, compared to increased levels of these transcripts by CIN micronuclei at 24 hours (Figure S6F-G). Overall, these data indicate that MSI micronuclei induce more effective STING activation at earlier time points in DCs, which has in other contexts been linked to more effective CD8^+^ T cell activation^16^.

### Lower levels of free cyDNA are required to induce similar levels of STING and CD8^+^ T cell activation to micronuclei

Next, we aimed to compare the relative ability of micronuclear vs free cyDNA to activate STING and induce CD8^+^ T cell activation by DCs. This is essential because studies throughout the literature either evaluate the role of cyDNA as a whole, investigate only micronuclei or free cyDNA, or only make comparisons to external DNA sources such as HT-DNA or interferon stimulatory DNA. Therefore, we isolated free cyDNA or micronuclei from MSI and CIN cells, used the DNA dye PicoGreen to normalize DNA input, and stimulated BMDCs with equal concentrations of these cyDNAs. Consistent with our previous findings^16^ (Figure 3E), free and micronuclear cyDNA from MSI cells led to increased expression of *Ifnβ* and *Cxcl10* compared to CIN cyDNA at early time points (Figure 4A). However, no consistent differences were observed in the expression of these genes between the free and micronuclear cyDNAs. As free cyDNA is known to contain ss and dsDNAs and PicoGreen stains only dsDNA, free cyDNA from MSI and CIN cells were treated with the ssDNA specific nuclease P1 before quantification by PicoGreen and stimulation of BMDCs. P1 nuclease treatment did not decrease *Ifnβ* expression induced by free cyDNA (Figure S7A), indicating that our results were not influenced by ssDNA in the MSI and CIN free cyDNA that were not quantified by PicoGreen, consistent with the known role of cGAS as a dsDNA sensor^15^.

**Figure 4.**
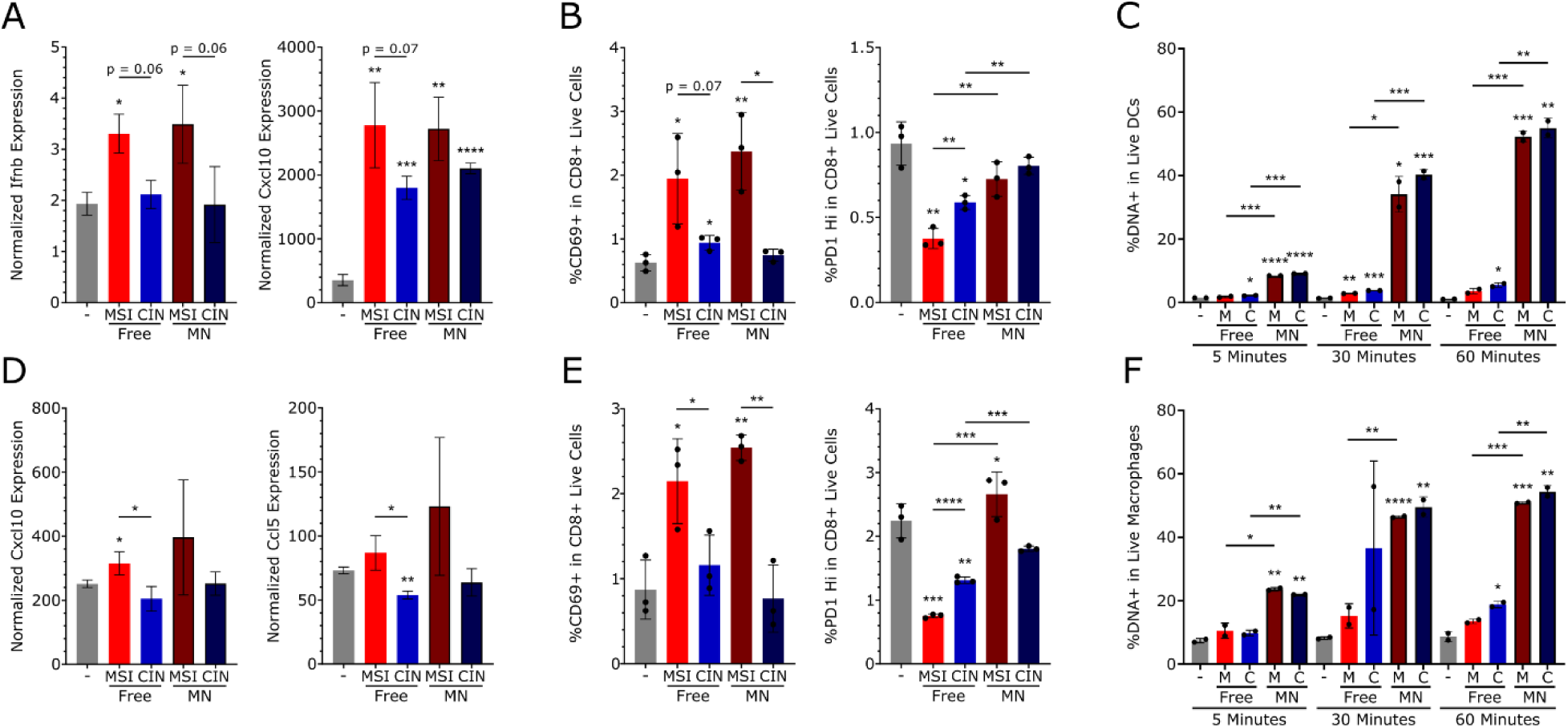
Less free cytosolic DNA is required for similar effects on STING and CD8^+^ T cell activation as micronuclei. (A, D) Micronuclei (MN) and free cyDNA were isolated from MSI and CIN cells, DNA concentration was normalized using PicoGreen quantification, and 500ng DNA was used to stimulate BMDCs for 3 hours (A) or BMDMs for 1 hour (D) before RNA isolation and RT-qPCR. Representative replicate shown of n = 3. (B, E) BMDCs (B) or BMDMs (E) were stimulated with 12.5ng micronuclei or free cyDNA from MSI and CIN cells for 30 minutes with OVA before washing and co-culture with OT-1 CD8^+^ T cells for 24 hours. Cells were then stained and T cell activation was measured by flow cytometry. Representative replicate shown from n = 3 with 3 biological replicates each. Data shown are normalized to T cell only controls. (C, F) MSI and CIN micronuclei and free cyDNA were pre-stained with PicoGreen before stimulation of BMDCs (C) or BMDMs (F) and evaluation of DNA uptake by flow cytometry. Representative replicate shown of n = 2. * over the sample bar indicates statistical significance to the lipofectamine vehicle control, all significance was determined by unpaired T test, * = p < 0.05, ** = p < 0.01, *** = p < 0.001, **** = p < 0.0001.

Next, to determine if micronuclear vs free cyDNA influenced activation of CD8^+^ T cells by DCs, MSI and CIN micronuclear and free cyDNAs were used with OVA to stimulate BMDCs for 30 minutes. After washing, stimulated BMDCs were then co-cultured with OT-1 CD8^+^ T cells for 24 hours before evaluation of CD8^+^ T cell activation by flow cytometry. Consistent with our previous observations^16^, equal concentrations of free MSI cyDNA led to greater upregulation of the T cell activation marker CD69 compared to free CIN cyDNA (Figure 4B). Likewise, MSI micronuclei stimulation improved CD69 levels on CD8^+^ T cells compared to CIN micronuclei. No differences in CD69 were observed between free and micronuclear cyDNAs. However, micronuclei from MSI and CIN cells led to more CD8^+^ T cells with high levels of the activation/exhaustion marker PD1. This is a critical observation since PD1 binding by its receptor PDL1 is a well described immune evasion strategy in tumors^45^, indicating micronuclei mediated PD1 upregulation may be important in establishing the cold tumor microenvironment (TME) in CIN CRCs.

Finally, although we utilized equal concentrations of free and micronuclear cyDNAs to stimulate the BMDCs, it is possible that our results could be influenced by differential uptake of these cyDNAs by DCs since this would lead to differences in the amount of cyDNA available in the cytosol to be sensed by cGAS. We therefore pre-stained MSI and CIN free and micronuclear cyDNA with PicoGreen before BMDC stimulation and analyzed DNA uptake using flow cytometry. Interestingly, approximately 10-fold more cyDNA was present in micronuclei stimulated BMDCs (Figure 4C, S7B). This effect did not appear to be due to longer DNAs in the micronuclei leading to more PicoGreen positivity per DNA strand, as similar results were observed with mean fluorescence intensity (MFI) and gating of percent positivity. Likewise, this could not be explained by differential staining efficiency by different DNA dyes as PicoGreen was used both for normalization of DNA input and quantification of uptake. Overall, these data suggest that far less free cyDNA from MSI or CIN cells is required to induce similar levels of cytokine expression and CD8^+^ T cell activation compared to micronuclear DNA.

### Micronuclei do not induce pro-tumorigenic inflammation in isolated immune cells

While we and others have shown many instances where STING improves anti-tumor immune responses leading to tumor prevention and treatment^9,17–19^, other studies have also linked STING activation to pro-tumorigenic inflammation and metastases^22,28^. It is not entirely understood how STING activation is modulated to achieve these different downstream effects, although one common theory postulates acute STING activation to be beneficial compared to chronic activation^46,47^. However, this theory contradicts observations that chronic STING activation from MSI cyDNA is beneficial to CD8^+^ T cell responses and patient prognosis^9^. Consistent with this, C57BL/6 mice pre-treated with the colitis inducer dextran sodium sulfate (DSS) before injection of MSI and CIN CRC cells into the colon show no differences in CD8^+^ or CD4^+^ T cell activation as measured by IFNγ (Figure S7C). Alternatively, we noticed a trend in the literature where pro-tumorigenic STING effects were frequently observed as a result of CIN and/or micronuclei^22,28^. We therefore asked whether micronuclei could sway STING activation towards pro-tumorigenic inflammation.

To investigate this, we first characterized the effects of free vs micronuclear cyDNA on STING activation in macrophages, whose phenotypic changes are reported to be modulated by STING and heavily influence pro- vs anti-tumor immunity^48^. Consistent with our observations in DCs, stimulation of unpolarized bone marrow derived macrophages (BMDMs) with equal concentrations of free or micronuclear cyDNA isolated from MSI and CIN cells showed no consistent differences in expression of *Cxcl10* or *Ccl5* between free or micronuclear cyDNAs (Figure 4D). Further, co-culture of OVA and free or micronuclear cyDNA stimulated BMDMs with OT-1 CD8^+^ T cells also led to similar levels of CD69 but increased PD1 in micronuclei treated conditions as was observed for DCs (Figure 4E). Likewise, BMDMs were also more effective at uptake of micronuclei cyDNA compared to free (Figure 4F, S7D). Therefore, as in DCs, far less free cyDNA is required for similar levels of cytokine expression and CD8^+^ T cell activation to micronuclei in macrophages.

We next determined whether free or micronuclear cyDNA influenced macrophage polarization. Stimulation of BMDMs with equal concentrations of free vs micronuclear cyDNA led to no consistent differences in expression of the M2 macrophage polarization markers *Arg1*, *Cd206*, or *Cd163* (Figure S7E), suggesting that micronuclei vs free cyDNAs do not alter classical macrophage polarization.

### Micronuclear cyDNA is less effective at inducing anti-tumor cytotoxic T cell responses in the TME

Next, to more comprehensively evaluate the role of free and micronuclear cyDNAs in anti- vs pro-tumor effects of STING, MSI and CIN cells were injected orthotopically into the colon wall of C57BL/6 mice (Figure 5A). We then treated these mice with BMDCs that had been stimulated with free or micronuclear DNA isolated from MSI CRC cells to determine if this could alter the TME. Tumors were then collected on day 15 for isolation of RNA and bulk RNA sequencing. Analysis of differentially expressed genes by gene ontology enrichment (GO) was used to determine pathways enriched by micronuclei or free cyDNA. Compared to micronuclei activated BMDCs, BMDCs stimulated with free cyDNA showed increased induction of several immune pathways in the TME including cytokine activity, T cell receptor complex, and response to bacterium in the tumor (Figure 5B). Comparatively, BMDCs stimulated with micronuclei led to increased T_H_17 cell regulation, negative regulation of T_H_ cell differentiation, and negative regulation of alpha-beta T cell differentiation (Figure 5C). At the level of specific genes, we found that free cyDNA treated DCs led to increased expression of *Cd8a*, *Acod1*, *MhcII*, and *Cd28* (Figure 5D). In contrast, micronuclei treated DCs led to increased expression of the Treg marker *Foxp3*, and the immuno-inhibitory cytokine *Il10* (Figure 5E). No difference was observed in the expression of *Il17a* (Figure S8A). These results demonstrate that while free cyDNA improves anti-tumor T cell responses by DCs, micronuclei instead lead to immunosuppression by DCs in CRC tumors.

**Figure 5.**
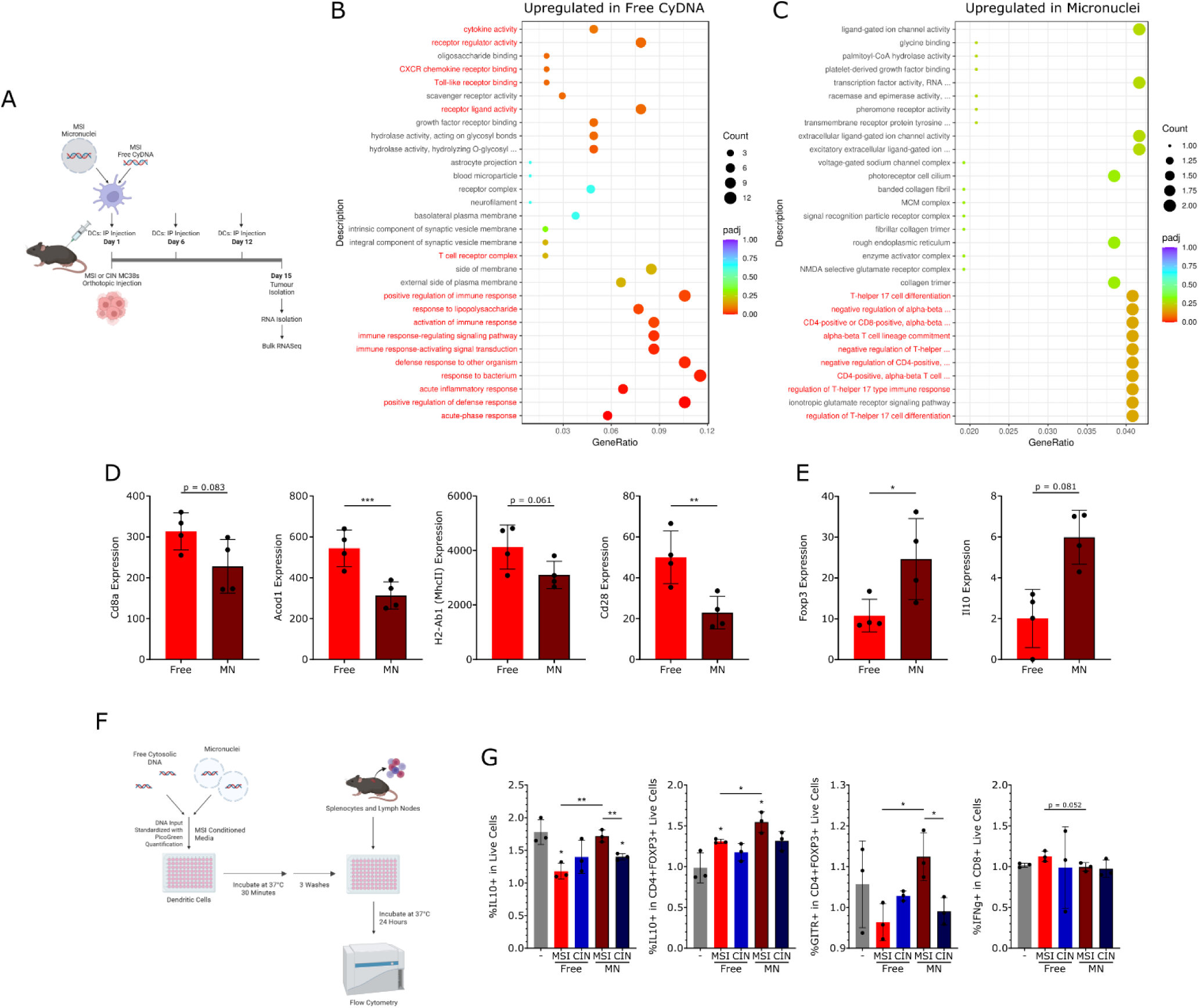
Free cyDNA leads to less IL-10 and more effective CD8^+^ T cell activation. (A-E) Free and micronuclear (MN) cyDNA was isolated from MSI cells and 2µg was used to stimulate 2×10^6^ BMDCs with lipofectamine. BMDCs were then injected intraperitoneally on days 1, 6, and 12 into C57BL/6 mice injected with 5×10^5^ MSI or CIN cells in the colon wall. On day 15, tumors were collected from which RNA was isolated and subjected to bulk RNA seq. Each point refers to RNA from 3 mice normalized and combined before RNA sequencing, n = 2 experimental replicates. (A) Experimental schematic, made in BioRender. (B-C) Dot plots of GO analysis of pathways upregulated in free compared to micronuclear cyDNA (B) or micronuclear compared to free cyDNA (C). Immune-related GO terms are coloured red. (D-E) Normalized read counts of the indicated genes upregulated in free (D) or micronuclear (E) samples. (F-G) 12.5ng free or micronuclear MSI cyDNA was used to stimulate 1×10^4^ BMDCs using lipofectamine with conditioned supernatant from MSI cells for 30 minutes. After washing, BMDCs were co-cultured for 24 hours with 1×10^5^ cells from the spleen and lymph nodes of C57BL/6 mice before evaluation of cytokine expression and CD4 and CD8 T cell activation by flow cytometry (G). Data shown is normalized to splenocyte only controls, n = 3 experimental replicates with 3 biological replicates each. Representative replicate shown. * over the data bar indicates significance to the lipofectamine vehicle control. Statistics determined by Wald Test (D) or unpaired T test (G), * = p < 0.05, ** = p < 0.01, *** = p < 0.001.

Given its important immunosuppressive role, we next aimed to examine the cell source and consequences of IL10 production induced in the TME by micronuclei compared to free cyDNA. Stimulation of BMDCs or BMDMs with micronuclei or free cyDNA isolated from MSI and CIN cells did not influence expression of *Il10* in the DCs or macrophages directly (Figure S8B-C), suggesting these cells instead regulate the ability of other cell types to produce IL10. Additionally, while previous studies reported IL10 production via STING-dependent induction of IDO, we observed no difference in *Ido1* expression in our RNA sequencing dataset (Figure S8D). To evaluate the role of micronuclei in regulating DC activation of other immune cell types in the TME, BMDCs were stimulated with equal concentrations of micronuclei or free cyDNA from MSI and CIN cells for 30 minutes in the presence of conditioned media from MSI cells (Figure 5F). After washing, BMDCs were then co-cultured with a mixed population of immune cells from the spleen and lymph nodes for 24 hours before examining IL10 production and immune composition by flow cytometry. Micronuclei from MSI cells led to increased production of IL10 in CD4^+^FOXP3^+^ Tregs as well as upregulation of the Treg activation marker GITR compared to free cyDNA from MSI cells (Figure 5G). Consistent with this, CD8^+^ T cell activation as measured by IFNγ was decreased in the MSI micronuclei condition compared to the free cyDNA. Overall, these data indicate that micronuclei lead to increased IL10 production by Tregs and are less effective at inducing CD8^+^ T cell responses by DCs. These findings show that the type of cyDNA present in the TME plays a critical role in determining the immunostimulatory vs immunosuppressive nature of MSI and CIN CRCs.

## Discussion

Understanding the unique features of MSI CRCs that enable them to mount effective anti-tumor immunity and respond to immune checkpoint inhibitors could provide critical information for how to induce similar favourable responses in CIN CRCs and other poorly immunogenic cancers. We previously showed that abundant production of CXCL10 and CCL5 by STING activation from highly stimulatory MSI cyDNA was an important factor in this process but did not address the critical question of what specific structural features in MSI cyDNA caused this. Here, we demonstrate that enrichment of G4s in the cyDNA from MSI CRCs are central to their potent antitumor immunity. Endogenous G4 structures in MSI cyDNA or in cyDNA of cells treated with a G4 stabilizer improved STING and CD8^+^ T cell activation by DCs. We additionally show that MSI micronuclei are more effective at inducing PRR activation than CIN micronuclei, however they were less effective at inducing CD8^+^ T cell activation than DCs treated with free MSI cyDNA. Critically, we found that micronuclear cyDNA led to more Treg activation and IL10 in the TME and thus could negatively impact overall anti-tumor immunity. Collectively, this work highlights the importance of understanding how different cyDNA structures regulate immune function and that therapies designed to activate the STING pathway must be carefully crafted to prevent undesired immune regulatory effects.

We are not the first to report G4s in the cytosol following DNA damage. However, to our knowledge we are the first to link G4 cyDNA to MSI and directly compare the efficiency of G4s compared to other cyDNAs in activating STING. Interestingly, a study by Byrd *et al*^49^ showed increased cytosolic G4s following oxidative damage with H_2_O_2_, which led to the development of stress granules. Of note, we observed that G4 high oligos did not show improved binding to cGAS, suggesting other mechanisms, such as possible impaired degradation by cytosolic nucleases, could be responsible. Likewise, while we did not observe improved ISG expression from the G4 high oligo B, this oligo significantly improved CD8^+^ T cell activation by DCs, suggesting that G4s may also be working through another PRR pathway such as the inflammasome and that further understanding of the role of specific G4 structures in immune regulation is required.

Previous reports have highlighted the ability of G4 binding compounds to induce STING activation and thereby improve cytotoxic T cell responses^43^. Our data further supports this by showing that increased G4s in the cytosol following treatment with G4 binding compounds leads to more effective STING activation, particularly in CIN CRCs. Cyclic dinucleotide STING agonist therapies have proven effective in sensitizing tumors to checkpoint inhibitors in animal models^50^, and further work is underway to improve STING agonist efficacy and limit toxicity, particularly through more effective delivery methods^51^. Early clinical trials showing limited toxicities using G4 stabilizing agents^30^ suggests these agents may prove an effective strategy of STING activation for combination with immune checkpoint blockade. Indeed, Chung *et al*^52^ observed combination of the G4 binder CX-5461 with either anti-PD1 or anti-PDL1 checkpoint therapies led to decreased tumor growth compared to either compound alone without loss of body weight. As G4s in the *KRAS* promoter, a gene upregulated in 40% of CRCs^2^, have also been shown to silence *KRAS* expression^42^, this combination of immune activation and oncogene silencing may be of particular interest in the treatment of CIN CRCs.

Current studies on STING activation often fail to separate the role of micronuclear vs free cyDNA or only evaluate the presence of micronuclei vs free cyDNA by fluorescent imaging at one time point without separating the possibly different timelines and roles of these cyDNA populations on downstream effects of STING. This has largely been due to a lack of tools to separate these two cyDNA types and difficulties in defining the specific differences between these cyDNA populations, particularly following micronuclear membrane rupture. Given that our investigation into the stimulatory nature of different cyDNA structures required separate isolation of free and micronuclear cyDNAs, we had a unique opportunity to directly compare the effects of these two cyDNA types on STING activation and anti-tumor immunity. Our observation that less free than micronuclear cyDNA was required to induce similar levels of cytokine expression in DCs was consistent with the findings of Takaki *et al*^44^ where micronuclei were less effective at inducing pSTING and IRF3 translocation compared to 94bp dsDNA oligos. These differences in STING activation may be due to the presence of nucleosomes on chromatin within micronuclei, which have been shown by several studies to inhibit cGAS^44,53,54^, or the presence of the micronuclear envelope, as previous work by Mackenzie *et al*^55^ highlights the requirement for micronuclear envelope rupture for cGAS sensing of micronuclear DNA. Further investigation will be required to determine which of these components influences differences in micronuclei vs free cyDNA recognition and the role of differential PRR activation by MSI and CIN micronuclei.

Additionally, we observed distinct downstream effects on anti-tumor immunity from micronuclei vs free cyDNA, with free cyDNA promoting CD8^+^ T cell activation and micronuclei promoting increased Treg activation and IL10 production. Notably, these differences were not evident in evaluations of STING activation in isolated DCs or CD8^+^ T cells, highlighting the importance of evaluating the effects of STING activation within the tumor microenvironment. While there have been reports of IL10 production by STING agonism^56,57^, these were observed as direct effects on the stimulated cell populations, whereas we did not observe consistent differences in IL10 expression directly in micronuclei stimulated DCs or macrophages. A study by Lemos *et al*^23^ also identified IL10 in tumor draining lymph nodes of Lewis lung carcinoma models that was dependent on *Sting* and *Ido1*, however we observed no differences in *Ido1* expression in micronuclei vs free stimulated tumors. In contrast, a study by Wilson *et al*^58^ found IL10 was induced by type I IFN during persistent viral infection. Further investigation will be required to determine if micronuclei induce IL10 and diminish CD8^+^ T cell responses through this mechanism. A recent study by Zhou et al^59^ found cyclic dinucleotide mediated STING activation in ILC3s increased intestinal Treg numbers and immune tolerance to the microbiome by increasing ILC3 homing to the mesenteric lymph node and antigen presentation. It is therefore interesting to consider whether micronuclei may induce DCs to activate Tregs directly or through other cell types, such as ILC3s. Overall, these data suggest that free and micronuclear cyDNA influence the balance between anti-tumor immunity and immunosuppression, where from a therapeutic standpoint it would be desirable to inhibit micronuclei effects and stimulate free cyDNA effects to optimize anti-tumor immune responses. Notably our investigations comparing micronuclei and free cyDNA only focused on untreated MSI and CIN micronuclei, which may be less effective at inducing anti-tumor immunity compared to micronuclei induced by DNA damaging therapies. Indeed, MacDonald *et al*^60^ found that cGAS recruitment to micronuclei varied according to the genotoxin used and identified histone modifications such as H3K79me2 that regulated this process.

Overall, our work identifies novel cyDNA structures produced by different forms of genomic instability that activate STING in DCs and thereby CD8^+^ T cells. We further identify a novel role for immunosuppression by STING following micronuclei stimulation. These results highlight the critical importance of understanding how different DNA structures differentially induce immune activation in the TME and must be carefully considered in the design of immunotherapies. Moving forward, our work will enable improved design of DNA-based STING agonists as a neoantigen independent immunotherapy that could help sensitize immunosuppressive tumors to current checkpoint inhibition therapies.

## Acknowledgements

The authors thank Dan McGinn, Cheryl Santos, Daming Li, Tatjana Golovin, Suey Van Baarle, Wayne Moffat, Jennifer Jones, Jeffrey Kwasney, Dr. Anne Galloway, and Dr. Lei Li, as well as the Analytical and Instrumentation and Cell Imaging cores at the University of Alberta for technical support. Dr. Avalyn Stanislaus and Dr. Michael Sawyer provided experimental support and advice. This project was supported by funding from the Canadian Institute of Health Research (grant 407882) (K.B.), the National Sciences and Engineering Research Council of Canada (grant RGPIN-2016-05152) (K.B.), the University Hospital Foundation (K.B.), the Cancer Research Institute of Northern Alberta (K.B.), a UKRI Future Leaders Fellowship (MR/W00738X/1) and the Henry Royce Institute for Advanced Materials, (EP/R00661X/1, EP/S019367/1, EP/P02470X/1 and EP/P025285/1) (A.L.B.P).

## Methods

### Cells

MC38 mouse colorectal cancer cells were rendered MSI (*ΔMlh1*) or CIN (*Kras^mut^*) by CRISPR induced mutation as previously reported^9^. *Mlh1* is silenced in the majority of sporadic MSI CRCs^5^ and activating *Kras* mutations are present in 40% of CRCs^2^. Mutated cells were cultured continuously for 6 months before use to allow accumulation of mutations consistent with each CRC subtype. MC38 cells were cultured in high glucose DMEM (10% FBS, 1% Penicillin/Streptomycin, 1% HEPES) at 37°C with 5% CO_2_.

For generation of BMDCs and BMDMs, femur and tibia bones were collected from C57BL/6 mice and bone marrow was flushed out with PBS. Cells were then incubated in ACK buffer (150mM ammonium chloride, 10mM potassium bicarbonate, 0.1mM EDTA, pH 7.4) for 2 minutes (BMDMs only) before a PBS wash. Cells were then plated in non-tissue culture treated plates at 5×10^6^ cells/8mL in BMDC (RPMI, 10% supernatant from B16-GMCSF cells, 5% FBS, 10mM HEPES, 1% Penicillin/Streptomycin, 50µM β-Mercaptoethanol) or BMDM (RPMI, 5µg/mL MCSF (StemCell), 10% FBS, 10mM HEPES, 1% Penicillin/Streptomycin, 50µM β-Mercaptoethanol) growth media. 8mL fresh media was added at days 3 and 5, and non-adherent (BMDC) or adherent (BMDM) cells were collected and frozen down for use on day 7. *Sting* and *Myd88* knockout BMDCs were generated from *Sting^Gt/Gt^* (JAX:017537) or *Myd88* (JAX:017537) knockout mice.

For isolation of splenocytes, the spleen and lymph nodes were collected from C57BL/6 mice. Spleens were strained through a 40µm filter before 5 minute incubation with ACK buffer. Lymph nodes were then strained through a 40µm filter and combined with spleen cells for use in co-culture experiments. For isolation of CD8^+^ T cells, spleen and lymph nodes were collected from OT-1 C57BL/6 mice (JAX:003831) and strained as above before negative selection using the EasySep Mouse CD8^+^ T cell Isolation Kit (StemCell) as per manufacturer instructions.

### Mice

C57BL/6 mice originally purchased from Charles River and C57BL/6 OT-1 mice originally purchased from Jackson Laboratory were bred and maintained at the Cross Cancer Institute vivarium. *Sting^Gt/Gt^* and *Myd88* knockout mice were purchased from Jackson Laboratory (JAX:017537, JAX:017537). Mixed groups of male and female littermates 8-20 weeks old were used for experiments. All animal work was approved by the Cross Cancer Institute’s Animal Care Committee.

### Free CyDNA isolation

Free cyDNA was isolated as previously described^61^. In brief, 5×10^6^ cells were collected by trypsinization before a PBS wash and resuspension in 550µL cytosolic extraction buffer (150mM NaCl, 50mM HEPES, 200µg/mL Digitonin (Millipore), 1M Hexylene Glycol (Fisher)) and incubation for 10 minutes on ice. Samples were then centrifuged at 2000 x g for 10 minutes at 4°C before collection of supernatant containing the cytosolic fraction. Cytosolic protein and RNA were then removed by incubation with Proteinase K (1mg/mL at 55°C for 1 hour, Fisher) and RNase A (500µg/mL at 37°C for 1 hour, Invitrogen), each of which was followed by phenol/chloroform/isoamyl alcohol extraction. This method was confirmed to have no detectable mitochondrial contamination by western blot for mitochondrial markers before removal of protein as shown previously^61^. All DNA damaging treatments used were confirmed to be non-lethal at the time of cyDNA isolation by crystal violet assay. For P1 nuclease treatment, the cytosolic fraction was incubated in 200U/mL P1 Nuclease (NEB) and 1X NEB Buffer 1.1 for 30 minutes at 37°C before the final phenol/chloroform/isoamyl alcohol extraction.

### Circular Dichroism

Free cytosolic DNA was isolated as above before separation by size exclusion chromatography using a HiLoad Superdex 75 10/300 column (Cytiva) pre-equilibrated with G4 buffer (20mM HEPES, 100mM KCl, 1mM EDTA, pH 7.5). For each run, 0.5mL of each sample was loaded onto the column and eluted at 0.65mL/min with G4 buffer. After collection of large and small cyDNA fractions (as shown in Figure S1A) and concentration using a 3kDa spin column (Cytiva), cyDNA was visualized from 220-320 nm on the Olis DSM 17 Circular Dichroism spectrometer using a cylindrical 1cm quartz cell. Data was analyzed with Olis Globalworks by blank subtraction, addition of a digital filter (11-13), and conversion to molar ellipticity. Circular dichroism analysis of G4 high and G4 low samples was preformed similarly without further separation by size exclusion chromatography.

### CyDNA sequencing

CyDNA isolated from MSI (2 clones of *ΔMlh1*), CIN (*Kras^mut^*, *ΔRad51*, CRISPR vector control), or *ΔPolε* MC38 cells were previously sequenced and published by us^16^ and deposited on the NCBI Sequence Read Archive: PRJNA1152723. Mapped and unmapped sequencing reads were filtered to remove bacterial contamination as previously described^16^ before evaluation of secondary structure using G4Hunter^32^, Gquad (https://cran.r-project.org/web/packages/gquad/index.html), and DNAShapeR^34^ packages. For DNAShapeR analysis requiring the same read sizes, reads were binned into 20bp size increments and trimmed equally off both edges to match the lowest bp size in the bin using BioPython SeqIO^62^. For improved visualization of means per base, curves were subjected to second order smoothing to the 4 closest neighbors using GraphPad Prism 8. All graphing was done with GraphPad Prism 8. Figures were combined in Inkscape.

### Atomic Force Microscopy (AFM)

Each sample was diluted to 3ng/uL in either 100mM KCl, 20mM HEPES pH 7.4 or 100mM LiCl, 20mM HEPES pH 7.4. From this dilution 2μL was immobilised (6ng final) on a freshly cleaved mica disk in 30μL of immobilization buffer (3mM NiCl2, 20mM HEPES, pH 7.4) for 5 min. The mica was then washed 4 times with 30μL of fresh immobilisation, before a further 30μL was added for imaging. All AFM measurements were performed in liquid following a previously published protocol^63^. All experiments were carried out in PeakForce Tapping imaging mode on a FastScan Dimension XR AFM system (Bruker), using FastScan D (Bruker) probes. The PeakForce amplitude was set to 10nm, the PeakForce Tapping frequency to 8kHKhz and the PeakForce setpoints in the range: 7-15mV, corresponding to peak forces of <70pN. Scans were taken with a size of 1×1μm at 1024 × 1024 pixels, at line rates of ∼3-5Hz. The freely available, open-source software TopoStats^64,65^ was used to process raw AFM data and analyse the DNA molecules (https://github.com/AFM-SPM/TopoStats), which is configured using a configuration file that can be found with the dataset. Briefly, the software loaded raw AFM images, carried out flattening, both line-by-line and plane flattening. Individual molecules were masked based on a height threshold to separate them from the background. A second flattening was carried out which excluded the grain, improving the flattening of the image. The height distribution of the flattened image was then shifted vertically to set the background to zero by calculating the mean of the non-grain containing data and subtracting that value from the image. Finally, a 1.1px Gaussian filter was applied to reduce any high-gain noise. Statistics on the grains were collated from “allstatistics.csv” and the volume of each were compared.

### BMDC cyDNA stimulation

BMDCs were thawed at 37°C for 3 minutes and resuspended in RPMI (10% FBS, 1% HEPES, 1% Penicillin/Streptomycin, 50µM β-Mercaptoethanol) before plating at 0.5×10^6^ cells per 2mL in 24 well plates. BMDCs were then stimulated with 500ng DNA and 0.125µL/mL Lipofectamine 2000 (Invitrogen) as per manufacturer instructions. For pulse stimulations (Figure 2E-F), BMDCs were seeded in 1mL RPMI in 2mL tubes before stimulation as above for 30 minutes. BMDCs were then washed with PBS before resuspension in RPMI and plating in 24 well plates for the remainder of the incubation time. Input concentrations were normalized by Nanodrop 1000 (Figure 2) or PicoGreen by: combining 4µL DNA and 100µL 1/200 PicoGreen (Thermo Fisher) in PBS, incubation at 37°C for 2 hours, and measurement on a microplate reader as per manufacturer instructions. Lipofectamine was not used to stimulate BMDCs in micronuclei stimulations, where PBS was used as a vehicle (Figures 3E-H). For stimulations to evaluate cyDNA uptake, free and micronuclear cyDNA were stained using 8µL/mL PicoGreen for 2 hours at 37°C before 50ng DNA (as determined by PicoGreen quantification above) was used to stimulate 5×10^4^ BMDCs or BMDMs with 0.125µL/mL Lipofectamine 2000 (Invitrogen) as per manufacturer instructions. 5 minutes before the end of the indicated incubation time, cells were spun down and resuspended in PBS for analysis using the CytoFlex S Flow Cytometer (Beckman). Cell viability was determined by DAPI stain. All western blot quantifications were performed using ImageJ.

### OT-1 CD8^+^ T cell co-cultures

BMDCs or BMDMs were thawed at 37°C for 3 minutes and resuspended in RPMI (10% FBS, 1% HEPES, 1% Penicillin/Streptomycin, 50µM β-Mercaptoethanol) before plating 5×10^4^ cells per 50µL in 96 well round bottom plates. BMDCs or BMDMs were then stimulated with 12.5ng DNA and 0.125µL/mL Lipofectamine 2000 (Invitrogen) as per manufacturer instructions with 0-5µg/mL OVA protein (Millipore Sigma). After 30 minutes at 37°C, cells were washed 3 times before addition of 1×10^5^ OT-1 CD8^+^ T cells and incubation at 37°C for the indicated time frame. Cells were then re-stimulated for 2 hours with 0.5µg/mL PMA (Millipore Sigma) and 50µg/mL Ionomycin (Thermo Fisher) before addition of 2µM Monensin (Thermo Fisher) and incubation for 2 more hours. Cells were then stained for flow cytometry using the listed antibodies (Table S3), Zombie Aqua Fixable Viability Kit (BioLegend), and the FOXP3/Transcription Factor Staining Kit (eBioscience) and run on the CytoFlex S Flow Cytometer (Beckman Coulter). Analysis was performed in FlowJo. For co-culture of BMDCs and splenocytes, the above protocol was performed with 1×10^4^ BMDCs and 1×10^5^ wildtype splenocytes. Instead of OVA protein, BMDCs were stimulated with DNA as described above in the presence of conditioned media from MSI CRC cells (3×10^6^ cells in 10mL of media for 24 hours). For visualization of IL10, cells were restimulated for 4 hours with 0.5µg/mL PMA (Millipore Sigma), 50µg/mL Ionomycin (Thermo Fisher), and 2µM Monensin (Thermo Fisher).

### Cytosolic G4 pulldown

50µL Protein G Dynabeads (Thermo Fisher) were washed 3 times in blocking buffer (0.5% BSA, PBS) before overnight incubation with 0.8µg BG4 antibody (Sigma) or IgG antibody (Thermo Fisher) in blocking buffer at 4°C on a rotisserie. Beads were then washed 3 times with blocking buffer and resuspended in 100µL blocking buffer. 8×10^6^ MC38 cells were collected by trypsinization and the cytosolic fraction was isolated by incubation in 500µL cytosolic extraction buffer (150mM NaCl, 50mM HEPES, 200µg/mL Digitonin (Millapore), 1M Hexylene Glycol (Fisher)) for 10 minutes on ice, spinning down at 2000 x g for 10 minutes at 4°C, and collection of the supernatant. This cytosolic fraction was then combined with the beads prepared above for incubation overnight at 4°C on the rotisserie. Before washing, supernatant was collected from the BG4 pulldown for isolation of G4 depleted cyDNA. Beads were then washed 6 times with wash buffer 1 (50mM HEPES, 500mM NaCl, 1mM EDTA, 1% NP40, 0.7% Na-deoxycholate) and twice with wash buffer 2 (50mM NaCl, TE). After resuspension of beads in elution buffer (50mM Tris-HCl pH 8.0, 10mM EDTA, 1% SDS), beads were incubated on a shaking heat block for 1 hour at 65°C before collection of supernatant containing immunoprecipitated cyDNA. Samples were then diluted 1:1 with TE, and all samples (including G4 depleted samples) were treated with RNase A (0.2mg/mL (Invitrogen), 37°C for 2 hours). 5mM CaCl_2_ was added, and samples were incubated with 0.2mg/mL Proteinase K (Fisher) at 55°C for 30 minutes. Following phenol/chloroform/isoamyl alcohol extraction, DNA was resuspended in G4 buffer (20mM HEPES, 100mM KCl, 1mM EDTA, pH 7.5) for further use.

### Preparation of synthetic G4s

Sequences of G4 oligos were designed by flanking a known G4 sequence with random sequence to a total of 70bp (Table S1). 5µM G4 oligo or scrambled control (IDT) was added to G4 buffer (20mM HEPES, 100mM KCl, 1mM EDTA, pH 7.5) and incubated for 10 minutes at 95°C and allowed to cool slowly overnight. G4 high and G4 low samples were then separated by size exclusion chromatography (Figure S5A) using a HiLoad Superdex 75 10/300 column (Cytiva) pre-equilibrated with G4 buffer. For each run, 0.5mL of each sample was loaded onto the column and eluted at 0.65mL/min with G4 buffer. Peak fractions were pooled and concentrated using a 3kDa spin column (Cytiva) for stimulation.

### Electrophoretic mobility shift assay (EMSA)

EMSA was performed as described in^35^. 1pmol of G4 high or low DNA oligo was combined with the indicated concentration of recombinant human cGAS protein (Cayman Chemicals) in EMSA buffer (150mM NaCl, 20mM HEPES) for a final volume of 10µL. After incubation for 30 minutes at room temperature, samples were kept on ice and 1µL of BlueJuice Gel Loading Buffer (Invitrogen) was added. Samples were then loaded onto a 2% agarose gel and run at 4°C in 0.5X TBE before staining in 1/10000 SYBR Gold (ThermoFisher Scientific) for 1 hour at room temperature on a rocker. Gels were visualized on the Amersham Typhoon (Cytiva) using the A^488^ channel.

### Micronuclei quantification

5×10^4^ MC38 CRC cells were plated in black 96 well plates (Greiner Bio-one) without or following treatment with 10Gy γ-IR. After 24 hour incubation, cells were fixed in ice cold methanol and incubated at -20°C for 15 minutes. Following 3 PBS washes, cells were incubated in blocking buffer (5% rat serum (StemCell), 0.3% Triton X-100, PBS) at room temperature for 1 hour before incubation with primary antibody (see Table S3) in antibody dilution buffer (1% BSA, 0.3% Triton X-100, PBS) for 1 hour. Cells were washed 3 times with PBS before secondary antibody incubation in antibody dilution buffer for 1.5 hours, 2 washes, incubation for 20 minutes with 1/100 Phalloidin (Invitrogen) in PBS, 2 washes, and incubation for 5 minutes with 1/2000 DAPI in PBS. Finally, cells were mounted in 70:30 glycerol:PBS and imaged on the HCS-ImageXpress XL microscope at 60X magnification (NA 0.85). Micronuclei analysis was performed with DAPI images on the Molecular Devices InCarta Image Analysis Software. Nuclei were identified by machine learning using the default model with the following modifications: area of 45-16468µm^2^, minimum intensity of 1500, and exclusion of objects touching the edge of the image. Micronuclei were identified by machine learning as trained on the images of replicate 1 with an area filter of 2-80µm^2^.

### Micronuclei isolation

Micronuclei isolation protocol was modified from^66^. Approximately 2×10^8^ cells were collected by trypsinization before resuspension in DMEM (10% FBS, 1% Penicillin/Streptomycin, 1% HEPES, 10µg/mL Cytochalasin D (StemCell)) and incubation at 37°C for 30 minutes. Cells were resuspended in 4mL Nuclei Extraction Buffer (Miltenyi Biotec) before transfer of 2mL each into 2 gentleMACs C Tubes (Miltenyi Biotec) and homogenization on the gentleMACS Octo Dissociator using the 4C_nuclei_1 protocol (Miltenyi Biotec). 4mL cell homogenate was then added to a sucrose gradient by layering 10mL 1.8M sucrose buffer (10mM Tris-HCl, 5mM magnesium acetate, 0.1mM EDTA, 1mM DTT, 0.3% BSA, 0.15mM spermine (Alfa Aesar), 0.75mM spermidine (Alfa Aesar), pH 8.0, with the indicated concentration of sucrose), 10mL 1.4M sucrose buffer, 10mL 1.06M sucrose buffer, then homogenate. Following centrifugation at 930 x g for 20 minutes at 4°C, the 1.8M fraction was collected and centrifuged at 35 000 x g for 90 minutes at 4°C. The pellet was then resuspended in 10mL 0.8M sucrose buffer, before layering onto a previously prepared linear gradient of 10mL 1.8M sucrose buffer and 10mL 1.0M sucrose buffer. Following centrifugation at 500 x g for 15 minutes at 4°C, 500µL fractions were collected from the top of the linear gradient. Fractions 2-6 contained micronuclei and were combined for further analysis. For nucleus bead purified micronuclei, the above protocol was performed along with bead purification using Anti-Nucleus Microbeads (Miltenyi Biotech) as per manufacturer instructions before continuing with sucrose gradient purification.

### Micronuclei electron microscopy

5µL of each micronuclei sample suspended in 0.1M HEPES was placed on a glow-discharged carbon film grid and negatively stained with 1% uranyl acetate for 1 minute before imaging with a 200kv JEOL 2100 Transmission Electron Microscope. For Figure 3D right, image was background subtracted to remove illumination gradient using ImageJ with a sliding paraboloid and rolling ball radius of 10,000 pixels.

### Micronuclei flow cytometry

For flow cytometry of micronuclei with PicoGreen, micronuclei were stained with 8µL/mL PicoGreen (Thermo Fisher) at 37°C for 2 hours before analysis on the Beckman CytoFlex S Flow Cytometer. For flow cytometry with cGAS, micronuclei were stained using the FOXP3 Transcription Factor Staining Kit (eBioscience) as per manufacturer instructions before PicoGreen stain as above.

### DSS Treatment

C57BL/6 mice were treated with 2% dextran sulfate sodium (DSS, Sigma-Aldrich) in the drinking water for 7 days, 14 days recovery with no DSS, then 2% DSS for 7 further days, and recovery for 3 days. 5×10^5^ MSI and CIN MC38 cells were injected orthotopically as performed previously using a flexible needle (Hamilton) inserted through the working channel of a Wolfe endoscope and visualized using the ColoView Imaging System^67^. After 19 days, tumors were collected, minced, and digested in a shaker for 30 minutes at 37°C in enzyme cocktail (RPMI, 10µg/mL DNaseI (Thermo Scientific), and 1mg/mL Collagenase IV (Sigma Aldrich). Tissue was then vigorously pipetted to dissociate cells and filtered through a 100µm strainer and washing. Cells were restimulated for 4 hours with 0.5µg/mL PMA, 50µg/mL Ionomycin, and for 2 hours with 2µM Monensin before flow cytometry staining with the indicated antibodies (Table S3), the Zombie Aqua Fixable Viability Kit (BioLegend), and the FOXP3/Transcription Factor Staining Kit (eBioscience). Stained samples were visualized on the CytoFlex S flow cytometer (Beckman Coulter). Analysis was performed in FlowJo.

### Bulk RNA sequencing

On day 1, C57BL/6 mice were injected orthotopically into the descending colon wall with 5×10^5^ MSI or CIN MC38 cells as performed previously using a flexible needle (Hamilton) inserted through the working channel of a Wolfe endoscope and visualized using the ColoView Imaging System^67^. Fresh BMDCs were prepared as described above and resuspended in RPMI at 2×10^6^ cells/mL for stimulation using 2µg MSI or CIN free cyDNA or micronuclei (as quantified using PicoGreen, Thermo Fisher) and 0.125µL/mL Lipofectamine 2000 (Invitrogen) per 2×10^6^ cells for 30 minutes at 37°C as per manufacturer instructions. Following 2 PBS washes, 2×10^6^ BMDCs were injected intraperitoneally into the tumor bearing mice on days 1, 6, and 12. On day 15, tumors were collected and digested using glass beads (Sigma Aldrich) and a Mini Beadbeater (Biospec Products) before RNA isolation using TRIzol (Invitrogen). Equal RNA quantities (as determined by the Qubit RNA BR Kit, Invitrogen) from each of the 3 biological replicates were then combined for bulk RNA sequencing performed by Novogene. Replicates shown (Figure 5A-E) are from 2 experimental replicates. Bioinformatic analysis was performed by Novogene. Gene expression data shown as FPKM normalized counts (Figure S8A) or normalized for sequencing depth (Figure 5D-E, S8D).

