## Supplemental Figures and Tables for "Cytosolic DNA structures produced by mismatch-repair deficiency coordinate anti-tumor immunity in colorectal cancer"

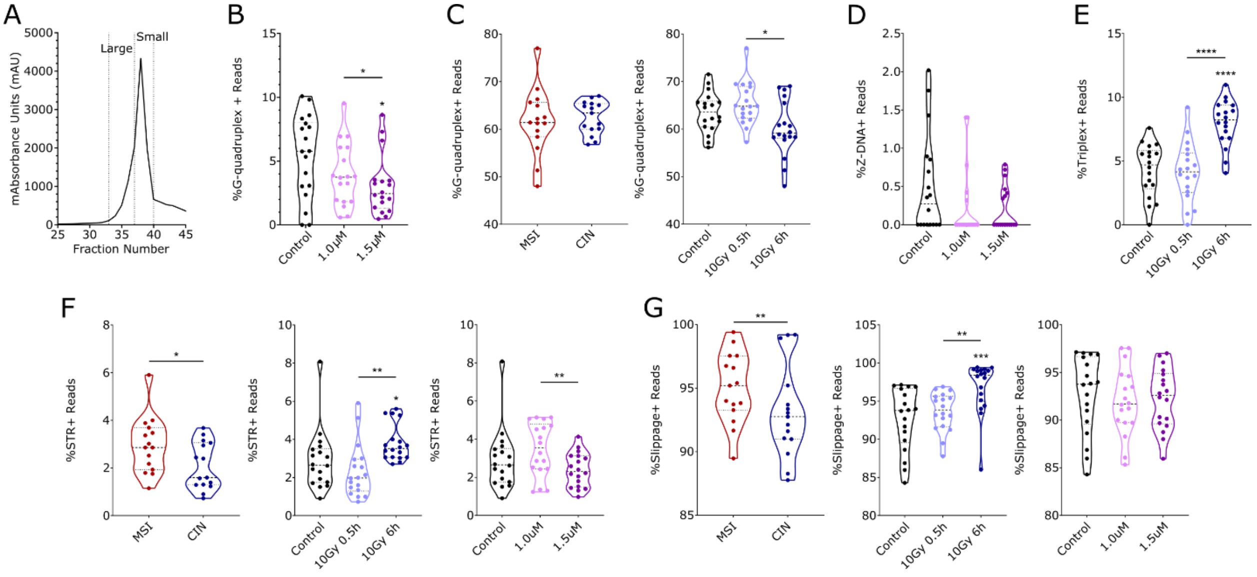

**Supplemental Figure 1. MSI and CIN cyDNA contain varying predicted DNA structures.** (A) CyDNA was isolated from MSI cells before separation by size exclusion chromatography into the indicated fractions for analysis by circular dichroism (Figure 1A) or AFM (Figure 1H-I, S4). (B-G) MSI and CIN cells were left untreated or treated with 10 Gy IR and allowed to recover for 0.5 or 6 hours or treated with 1.0 or 1.5µM 5-FU for 24 hours before cyDNA isolation and analysis of cyDNA content by next generation sequencing. See Figure 1B-G. (B) G4Hunter analysis of predicted G4 prevalence in CRC cells treated with 5-FU. (C) Gquad analysis of predicted G4 prevalence in MSI and CIN cyDNA (left) or CRC cyDNA after IR treatment (right). (D) Percent of reads that contain sequences that could form Z-DNA in the appropriate conditions in cyDNA from CRC cells treated with 5-FU. (E) Percent of reads containing predicted triplexes in cyDNA from CRC cells treated with IR. (F) Percent of reads containing small tandem repeats (STRs) in MSI vs CIN cyDNA (left), cyDNA isolated after IR treatment (middle), or cyDNA isolated after 5-FU (right). (G) Percent of reads containing slipped motifs in MSI vs CIN cyDNA (left), cyDNA isolated after IR treatment (middle), or cyDNA isolated after 5-FU treatment (right). Graphs shown contain all treatment groups within each cell type (C left, F left, G left), or all cell types within each treatment group (B, C right, D-E, F-G middle and right). Statistics were determined by paired T test, * above the data group indicates significance to the untreated control, * = $p < 0.05$, ** = $p < 0.01$, *** = $p < 0.001$.

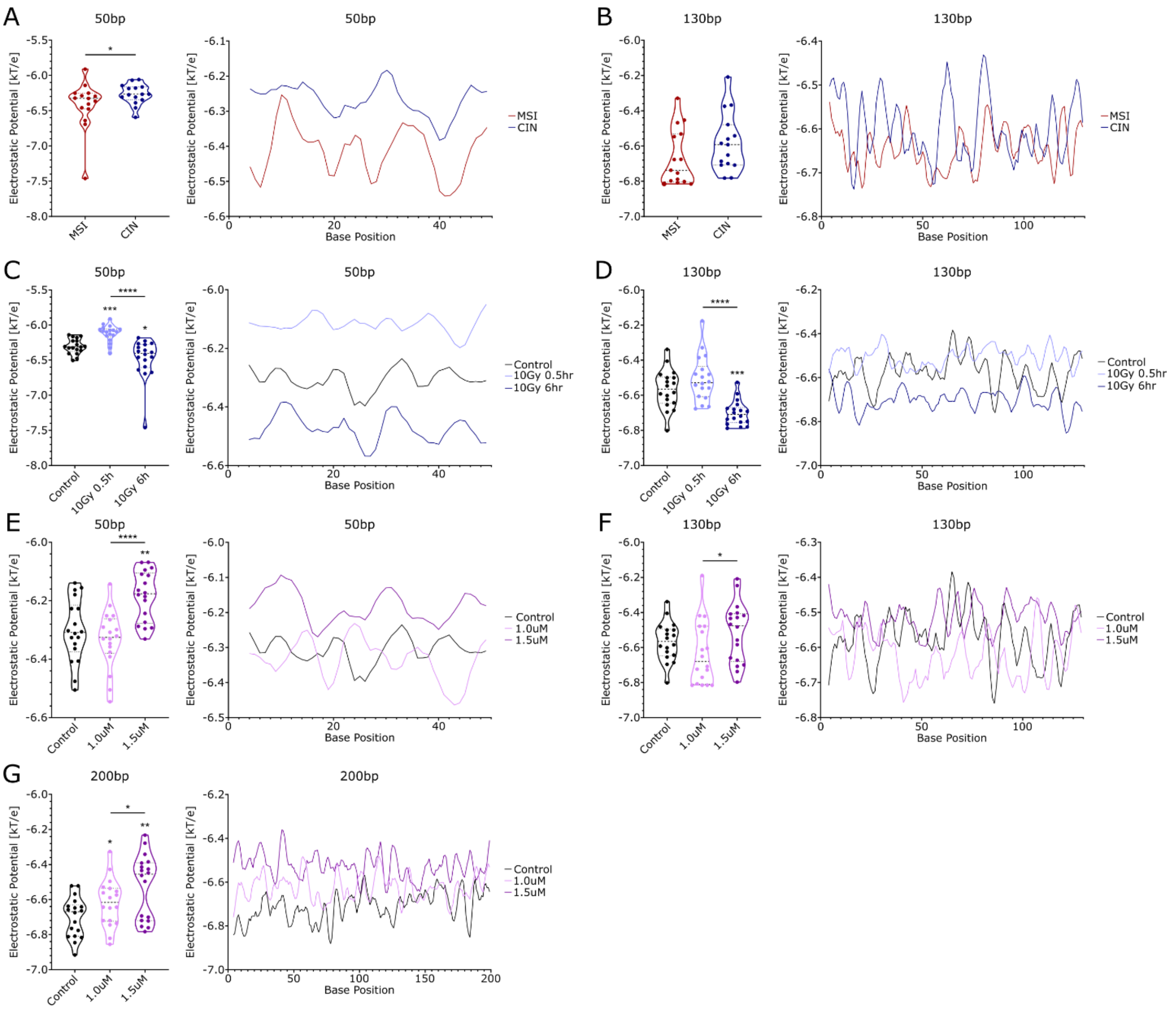


**Supplemental Figure 2. CyDNA electrostatic potential varies with CRC subtype and DNA damaging therapy.** MSI and CIN CRC cells were left untreated, were treated with 10 Gy IR and allowed to recover for 0.5 or 6 hours, or were treated with 1.0 or 1.5µM 5-FU for 24 hours before cyDNA isolation and sequencing. The DNAShapeR package was used to evaluate electrostatic potential in cyDNA sequences of each size range for MSI vs CIN (A-B), cyDNA after IR treatment (C-D), and cyDNA after 5-FU treatment (E-G). Mean per sample is shown on the left and mean per base is shown on the right. Data shown is all treatment types combined (A-B) or all CRC subtypes combined (C-G). Statistical significance was determined by paired T test, * over the sample values indicates significance to the untreated control, * = $p < 0.05$, ** = $p < 0.01$, *** = $p < 0.001$, **** = $p < 0.0001$.

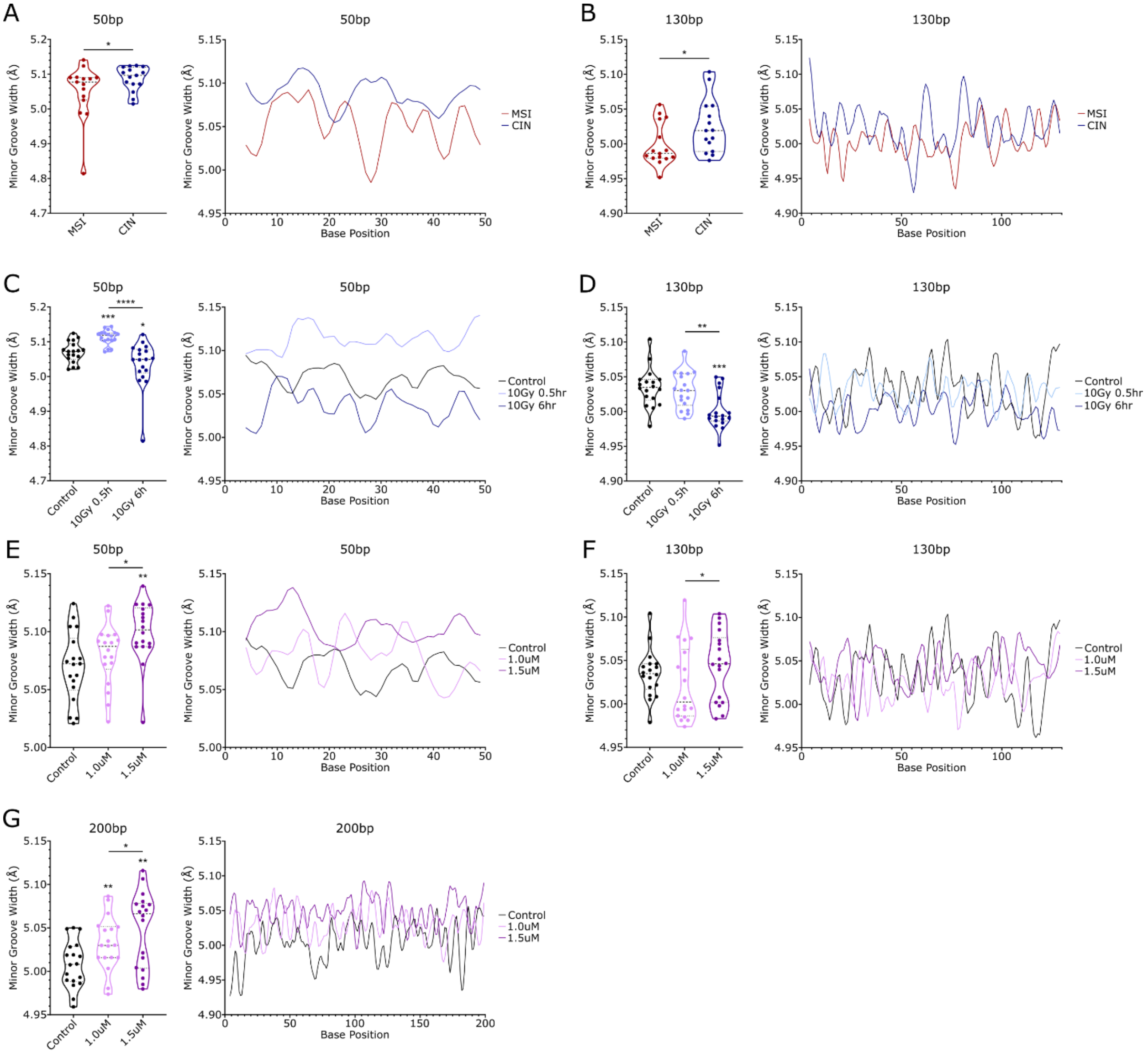

**Supplemental Figure 3. CyDNA minor groove width varies with CRC subtype and DNA damaging therapy.** MSI and CIN CRC cells were left untreated, were treated with 10 Gy IR and allowed to recover for 0.5 or 6 hours, or were treated with 1.0 or 1.5µM 5-FU for 24 hours before cyDNA isolation and sequencing. The DNAShapeR package was used to evaluate minor groove width in cyDNA sequences of the indicated size for MSI vs CIN (A-B), cyDNA after IR treatment (C-D), and cyDNA after 5-FU treatment (E-G). Mean per sample is shown on the left and mean per base is shown on the right. Data shown is all treatment types combined (A-B) or all CRC subtypes combined (C-G). Statistical significance was determined by paired T test, * over the sample values indicates significance to the untreated control, * = $p < 0.05$, ** = $p < 0.01$, *** = $p < 0.001$, **** = $p < 0.0001$.

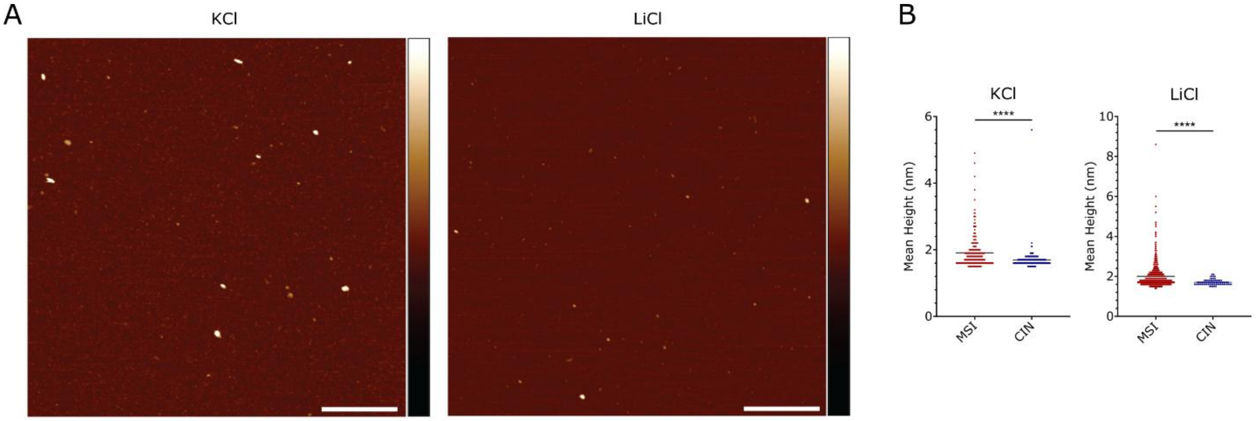


**Supplemental Figure 4. Large cyDNA fragments do not contain G4s.** MSI and CIN cyDNA was separated by size exclusion chromatography before imaging on the AFM. (A) Representative uncropped images of cyDNA in the approximately 40bp size range in KCl (left) or LiCl (right) buffers, see Figure 1H-I. Scale bars are 200nm and height scale bar is -3 to 4nm. (B) Mean height of each DNA strand in the >45bp size range in KCl (left) or LiCl (right) buffers. Statistical significance was determined by unpaired T test, **** = $p < 0.0001$.

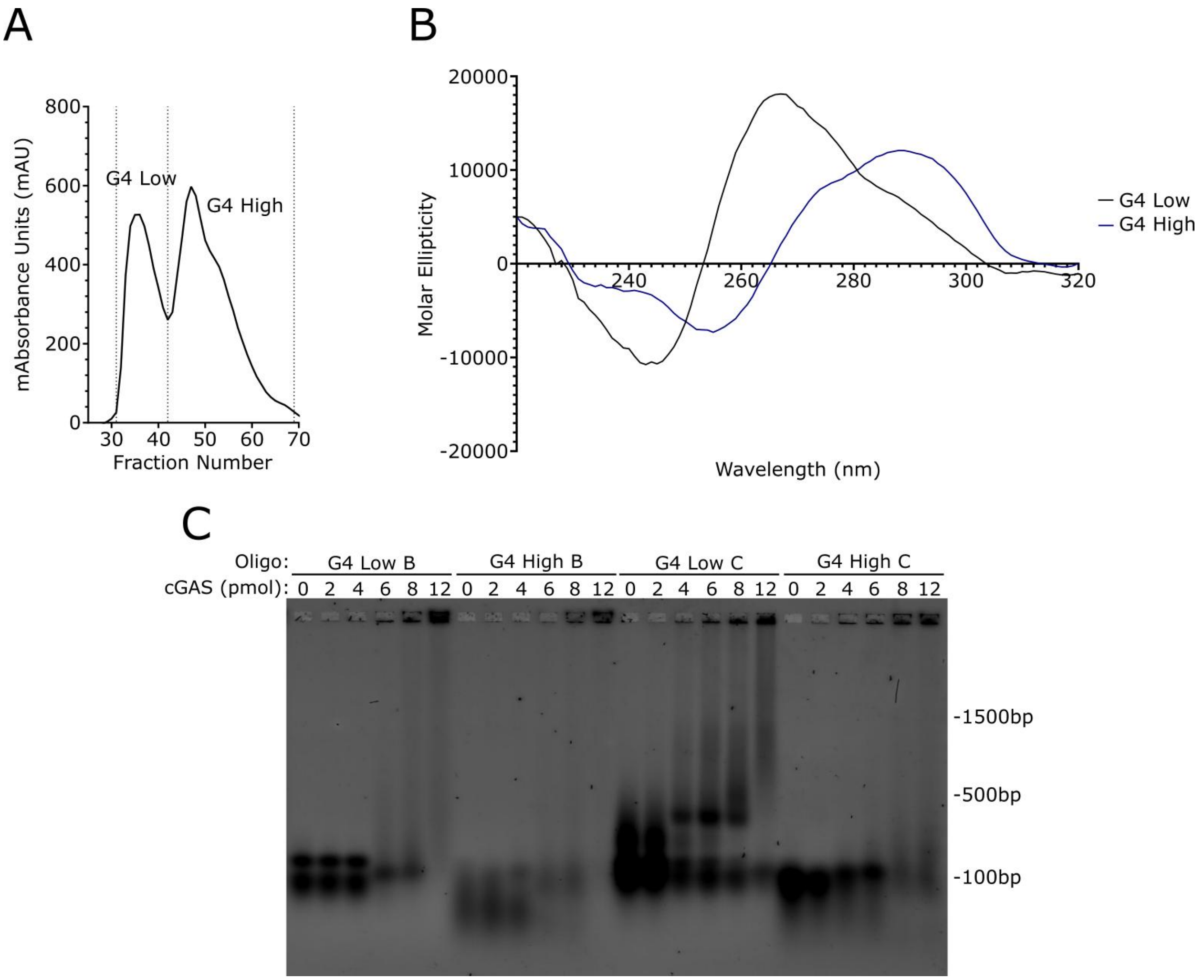


**Supplemental Figure 5. G4 high oligos do not exhibit improved cGAS binding to G4 low oligos.** Oligos containing sequences of known promoter G4s were allowed to form structure by heating and cooling slowly overnight before separation of oligos that formed structure (G4 high) and oligos that formed less structure (G4 low) by size exclusion chromatography for use in stimulations. (A) Fractions collected by size exclusion chromatography for G4 high and low oligo C. (B) Circular dichroism spectroscopy of G4 high and G4 low oligo C. (C) EMSA of cGAS binding to G4 high and low oligos B and C. Representative replicate shown of n = 2.

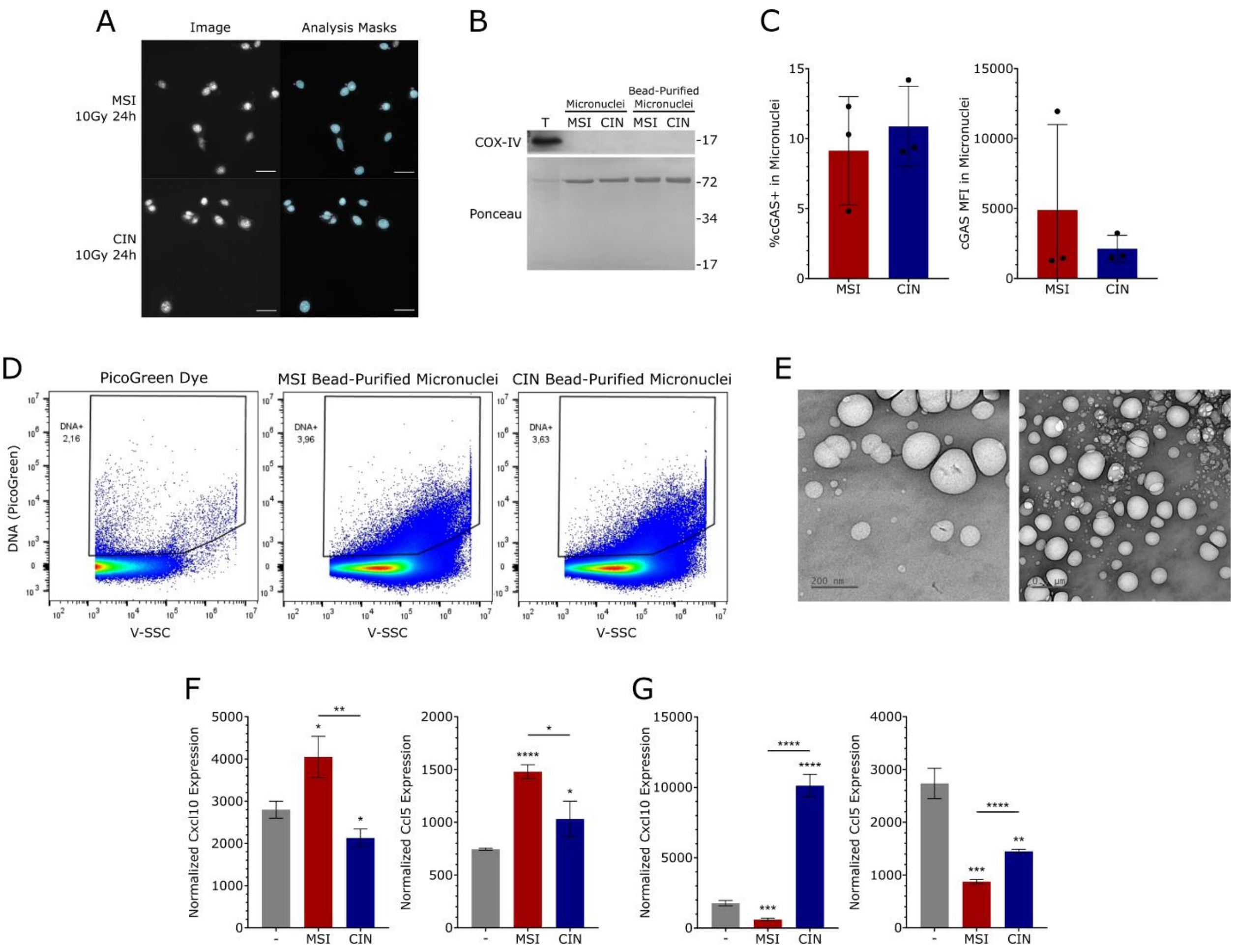


**Supplemental Figure 6. Micronuclei produced by MSI lead to more efficient PRR activation than CIN.** (A) Representative analysis masks for data shown in Figure 3A. MSI and CIN cells were treated with 10Gy IR, allowed to recover for 24 hours, and were fixed and stained with DAPI to evaluate micronuclei content. Blue indicates the nuclei counted, purple indicates the micronuclei counted. Scale bars are equal to 10µm. (B) Western blot of the mitochondrial marker COX-IV on micronuclei and bead purified micronuclei isolated from MSI or CIN cells. T indicates protein from total CIN cells. (C) Micronuclei isolated from MSI and CIN cells were stained with cGAS antibody and PicoGreen for micronuclei identification before analysis of cGAS content by microparticle flow cytometry. Data shown is combined from n = 3. (D) Bead purified micronuclei from MSI and CIN cells were stained with PicoGreen and DNA content was evaluated by microparticle flow cytometry. Representative replicate of n = 3. Flow graphs taken from FlowJo. (E) Bead purified micronuclei from CIN cells were imaged by Transmission Electron Microscopy. Scale bars are equal to 200 nm (left) or 0.5µm (right). (F-G) Bead purified micronuclei from MSI and CIN cells were normalized by DNA concentration using PicoGreen quantification and 100ng was used to stimulate BMDCs for 1 (F left), 3 (F right, G left), or 24 hours (G right). Representative replicate shown of n = 3. * over the sample bar indicates significance to the vehicle control. Statistical significance was determined by unpaired T test, * = $p < 0.05$, ** = $p < 0.01$, *** = $p < 0.001$, **** = $p < 0.0001$. * over the sample bar indicates significance to the vehicle control.

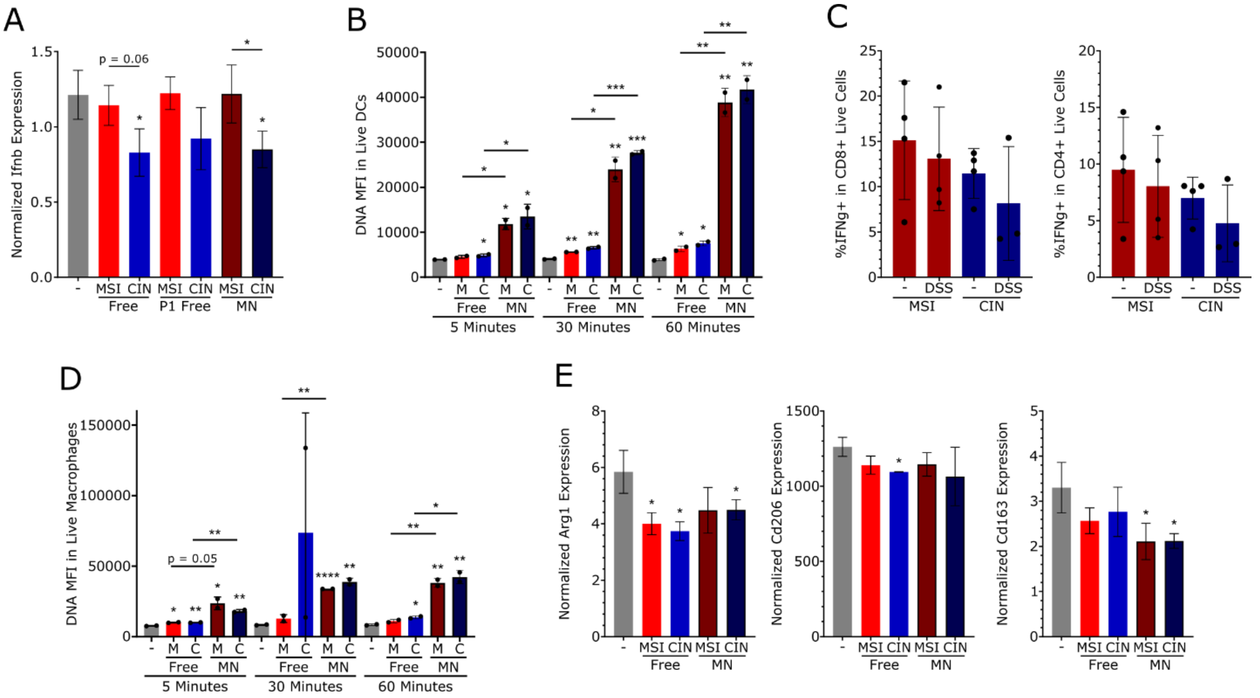


**Supplemental Figure 7. Free vs micronuclear cyDNA doesn't alter macrophage polarization.** (A) Free, P1 nuclease treated free, or micronuclear (MN) cyDNA from MSI and CIN cells were used to stimulate BMDCs for 1 hour before isolation of RNA and qPCR. Representative replicate shown of n = 2. (B, D) Free and micronuclear cyDNA were isolated from MSI and CIN cells, pre-stained with PicoGreen, and used to stimulate BMDCs (B) or BMDMs (D) for the indicated time frame before evaluation of cyDNA uptake by flow cytometry. Data shown is MFI in live cells. Representative replicate shown of n = 2. (C) C57BL/6 mice were pretreated with DSS (2% in the drinking water for 7 days, 14 days recovery, 2% in the drinking water for 7 days, 3 days recovery) before orthotopic injection of MSI or CIN cells into the colon wall. Tumors were allowed to grow for 19 days before tumor isolation and evaluation of immune cell composition by flow cytometry. Representative replicate shown of n = 3 experiments of 4 mice each. (E) BMDMs were stimulated with free or micronuclear cyDNA from MSI or CIN cells for 4 hours before RNA isolation and RT-qPCR. Representative replicate shown of n = 3. * over the sample bar indicates significance to the vehicle control. Statistical analysis was performed by unpaired T test, * = $p < 0.05$, ** = $p < 0.01$, *** = $p < 0.001$, **** = $p < 0.0001$.

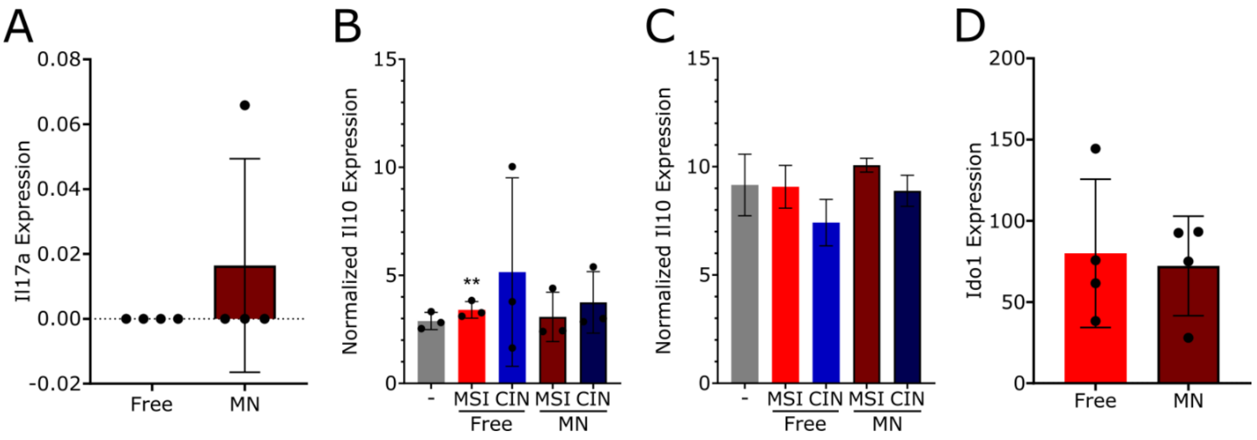


**Supplemental Figure 8. IL10 is not produced by DCs or macrophages directly stimulated with micronuclear or free cyDNA.** (A, D) See Figure 5A-E. Micronuclei or free cyDNA isolated from MSI CRC cells was used to stimulate BMDCs at equal concentrations before injection of BMDCs intraperitoneally into C57BL/6 mice bearing orthotopic CIN or MSI tumors. On day 15 tumors were collected, RNA was isolated, and bulk RNA sequencing was performed. RNA from tumors from 3 mice was normalized by RNA quantity and combined before sequencing. Data shown as FPKM normalized read counts, n = 2 experimental replicates. (B-C) BMDCs or BMDMs were stimulated with micronuclear (MN) or free cyDNA isolated from MSI or CIN cells before isolation of RNA and RT-qPCR. (B) BMDCs stimulated for 3 hours, each data point is an experimental replicate of n = 3. (C) BMDMs stimulated for 4 hours, representative replicate shown of n = 3. (D) See Figure 5A-E, S8A. Data shown as normalized read counts, n = 2 experimental replicates with 3 mice each. Statistical significance was determined by paired (B) or unpaired (C) T test, ** = $p < 0.01$.

**Supplemental Table 1. Sequences of DNA oligos.**

| Oligo | Figures Used | Sequence (5' to 3') |
|---|---|---|
| Scramble Fwd | Figure 2B-D | CTCCGGCGCCAAAGAGTATTGGGTCCTATCCACTGGTCAGCAAACTGTGCCATATGAAGCAAGTAACACG |
| Scramble Rev Complement | Figure 2B-D | CGTGTTACTTGCTTCATATGGCACAGTTTGCTGACCAGTGGATAGGACCCAATACTCTTTGGCGCCGGAG |
| G4 Oligo A | Figure 2B-D | TGCATGAAATGTTGCCGCGTCATCGGGGCGGGGCGGGGGCGGGGGCGACACAGTTGGTCGCATGCGCTTA |
| G4 Oligo B | Figure 2B-D | CACAGCTGCGCATACAGTAACGGGTTGCGGGCGCAGGGCACGGGCGGCGGTGGATGCCATAGGCCGTATA |
| G4 Oligo C | Figure 2B-D, S5A-B | GTGCTGCATGAATGTTCGTGGCTGGGGGCTCGGGGCGCGGGGCGCGGGGCATGGGGCATCGCCATCTGCA |

**Supplemental Table 2. Primers used for RT-qPCR.**

| Target | Sequence |
|---|---|
| *Gapdh* Fwd | CATGTTCCAGTATGACTCCA |
| *Gapdh* Rev | TGAAGACACCAGTAGACTCC |
| *Cxcl10* Fwd | CCAAGTGCTGCCGTCATTTTC |
| *Cxcl10* Rev | GGCTCGCAGGGATGATTTCAA |
| *Ccl5* Fwd | GCTGCTTTGCCTACCTCTCC |
| *Ccl5* Rev | TCGAGTGACAAACACGACTGC |
| *Irf7* Fwd | GGTGTGTCCCCAGGATCATT |
| *Irf7* Rev | GCTGCATAGGGTTCCTCGTAA |
| *Isg15* Fwd | GGTGTCCGTGACTAACTCCAT |
| *Isg15* Rev | TGGAAAGGGTAAGACCGTCCT |
| *Ifnβ* Fwd | CGTGGGAGATGTCCTCAACT |
| *Ifnβ* Rev | AGATCTCTGCTCGGACCACC |
| *Arg1* Fwd | CTCCAAGCCAAAGTCCTTAGAG |
| *Arg1* Rev | AGGAGCTGTCATTAGGGACATC |
| *Cd206* Fwd | TCATTCCCTCAGCAAGCGAT |
| *Cd206* Rev | CTGTCCGCCCAGTATCCATC |
| *Cd163* Fwd | CACGGCACTCTTGGTTTGTG |
| *Cd163* Rev | CTCTGAATGACCCCCGAGGA |
| *Il10* Fwd | GCTCTTACTGACTGGCATGAG |
| *Il10* Rev | CGCAGCTCTAGGAGCATGTG |

**Supplemental Table 3. List of antibodies used.**

| Target | Source | Identifier |
|---|---|---|
| G4 (BG4) | Sigma Millipore | MABE917 |
| Ki67 PECy5 | eBioscience | 15-5698-82 |
| CD8 FITC | eBioscience | 11-0081-82 |
| CCR5 APC | eBioscience | 17-1951-82 |
| pTBK1 | Cell Signaling Technology | 5483S |
| TBK1 | Cell Signaling Technology | 3504S |
| pSTAT1 | Thermo Fisher | 333400 |
| STAT1 | Cell Signaling Technology | 9172S |
| pNF-κB p65 | Cell Signaling Technology | 3033S |
| GAPDH | Thermo Fisher | PIMA515738 |
| cGAS | Cell Signaling Technology | 31659 |
| Anti-Rabbit $A^{647}$ | Jackson Immunoresearch | 111-606-144 |
| CD69 A700 | eBioscience | 56-0691-82 |
| PD1 SB600 | eBioscience | 63-9981-82 |
| IL10 PE | eBioscience | 12-7101-82 |
| GITR PECy7 | eBioscience | 25-5874-82 |
| IFNγ PE | eBioscience | 12-7311-82 |
| CD4 FITC | eBioscience | 11-0041-85 |
| FOXP3 APC | eBioscience | 17-5773-82 |
